# Angel Wing: a class of alternative splicing regulators in animals

**DOI:** 10.1101/2025.04.20.649734

**Authors:** Ting Zhang, Kaili Li, Ying Xiao, Xionghong Tan, Xiaoyan Tan, Min Cui, Yikang S. Rong

## Abstract

Alternative splicing (AS) regulates the diversity and level of the proteome. The specificity in AS is in turn regulated by RNA binding proteins, but our understanding of how they act is far from complete. Here we identify the Angel wing (Anw) protein as a novel AS regulator. Loss of Anw in *Drosophila* disrupts splicing in muscle genes and subsequently muscle function. Based on a mini-gene assay in which Anw and its RNA targets are co-expressed in cultured cells, we demonstrated orthologous splicing regulation of the minigene transcripts, interaction between Anw and its RNA targets, and a remarkable functional conservation among Anw homologs. Anw forms nuclear foci, and genetic ablation of Anw domains suggests that maintaining distinctive features of these foci is important for its function. The evolution of Anw is dynamic with gene gains and losses, but preserves a cross-phyla “ultra conserved element” as an alternative exon that potentially regulates its own level by non-sense mediated mRNA decay. As the human *anw* homolog is a candidate gene for myasthenia gravis, our work suggests a mechanism for cellular dysfunction in this disease.

## Introduction

RNA splicing, the process of removing intronic and joining exonic sequences, is an essential step of mRNA maturation in eukaryotes. The pattern of pre-mRNA splicing need not deterministic but allows alternative demarcation of exons. This process of Alternative Splicing (AS) increases the diversity of the eukaryotic proteomes by rendering a single pre-mRNA species capable of encoding more than one but sometimes thousands of protein isoforms. These isoforms often differ slightly in their primary sequences allowing them to fulfill molecular functions specially tuned for different tissues and developmental programs. Some of the best examples of this functional diversification lie in the muscle proteomes (e.g., Venables et al. 2012; Nikonova et al. 2020). The extent of AS appears correlated with organismal complexity (e.g., Soller 2006; Lee and Rio 2015), hence offering a potential explanation for why the number of genes in eukaryotic organisms correlates poorly with their organismal complexity. With about 20000 protein-encoding genes, the human genome has a genic transcriptome with one of the highest levels of AS at over 90% (Pan et al. 2008; Wang et al. 2008). In addition, the extent of AS appears the highest in complex tissues such as the muscular and nervous systems (Furlanis and Scheiffele 2018; Hinkle et al. 2019; Nikonova et al. 2020; Lee et al. 2023; Bao et al. 2024). Not surprisingly, many neuro- and muscular-diseases are rooted in defective AS.

AS can also serve as a regulatory mechanism for the level of individual proteins when coupled with the process of nonsense-mediated mRNA decay (NMD), which degrades mRNAs with premature stop codons. For example, some of the alternative exons harbor in-frame stop codons. These “poison exons” when included generate mRNAs that are degraded by the NMD machinery, thus serving to regulate the level of target proteins (reviewed in Lareau et al. 2007b; García-Moreno et al. 2020; Titus et al. 2021). Remarkably, some of the splicing factors themselves possess highly conserved poison exons, and use AS-NMD as a self-regulatory mechanism (Saltzman et al. 2008; Lareau and Brenner 2015; Li et al. 2024).

The complex AS patterns described for many animal species present a daunting task for the understanding of their functions (Graveley 2001). In particular, what portion of these AS events are noise or the results of splicing errors remains rather obscure. One way to identify AS events with functional significance is through loss of function studies. For example, circular RNA produced by the process of “back splicing” has been categorized as the result of splicing errors that confers no fitness benefit (Xu and Zhang 2021). The revelation that a small number of these circular RNAs do carry important functions has only become apparent after painstaking studies of the effects upon their loss (e.g., Pamudurti et al. 2022; Giusti et al. 2024). The fruit fly *Drosophila melanogaster*, with a similar number of genes as humans, but a simpler physiology or development, is one of the best models for conducting loss of function studies of AS, enhanced by one of the best categorized AS programs through development (Daines et al. 2011; Graveley et al. 2011).

In AS, different subsets of splice sites are used in cells of different tissues or developmental stages. This specificity is not based solely on primary RNA sequences of the splice sites as their defining features are rather rudimentary. Although secondary RNA structures formed within the alternatively spliced regions are often an additional landmark critical for AS specificity (e.g., Graveley 2005; Yang et al. 2011; Li et al. 2024), a more dominant role may be played by RNA interacting proteins that bind to the vicinity of the target splice sites and influence their selection. There are more than 200 RNA binding proteins (RBPs) annotated in *Drosophila* (Lasko 2000; Gamberi et al. 2006). The predominant class of fly RBPs is the 117 members that contain one and sometimes multiple copies of the domain called RNA recognition motif (RRM), a domain best characterized for its sequence specific RNA binding (Kenan et al. 1991; Maris et al. 2005). There are likely twice as many RRM-containing proteins encoded by the human genome (Gerstberger et al. 2014), a significant portion of which remain uncharacterized.

Traditionally, the role of RBPs in AS regulation was revealed in extensive characterization of the RBP mutants. One of the best examples is RBPs in regulating the cascade of AS events that ultimately determine the sexes in *Drosophila* (reviewed in Lopez 1998; Soller 2006; Venables et al. 2012). In a second example, the identification of ELAV/Hu as a class of RBPs essential in AS started from mutations causing defective neuronal development in embryos (e.g., Koushika et al. 1996; Soller and White 2005; Wei et al. 2020). Last but not least, in human myotonic dystrophy type 1 (DM1) patients, an aberrantly expressed RNA repeats results in the sequestration of the Muscleblind and Muscleblind-like (MBNL) proteins, leading to the discovery of the MBNL class of RBPs being important AS regulators (reviewed in Konieczny et al. 2014).

Through the study of RBPs’ role in AS, we gained much knowledge about commonly employed mechanisms (reviewed in Lopez 1998; Soller 2006; Park and Graveley 2007; Venables et al. 2012; Lee and Rio 2015; Ule and Blencowe 2019; Gehring and Roignant 2021; Lee et al. 2023; Bao et al. 2024). For example, RBP binding facilitates/blocks the recruitment of spliceosomal subcomplexes, leading to the inclusion/exclusion of alternative exons. RBP binding can also mediate long range interaction resulting in RNA looping that modulates splice site selection (Blanchette et al. 2009). Interestingly, nucleated binding of RBPs followed by their spreading over a large region can also modulate accessibility region-wide (Blanchette et al. 2009). More recently, a phase-separated multi-protein complex called the large assembly of splicing regulators has been identified in which cooperation amongst individual factors leads to different AS outcomes (Damianov et al. 2016).

The efforts are ongoing to find new RBPs important for AS, including both RNAi-based loss-of-function studies on Drosophila RBPs (Park et al. 2004; Brooks et al. 2015), and a more recent gain-of-function approach identifying human RBPs capable of mediating alternative exon inclusion (Schmok et al. 2024). Here we described the initial characterization of the founding member of a class of AS regulators, the Angel wing protein (Anw). Loss of Anw in *Drosophila* leads to flightless adults. In these animals, disrupted muscle organization is accompanied by altered AS programs involving multiple genes enriched with muscle functions. Anw protein has an RRM domain and the ability to form nuclear bodies. Interestingly, altering the number or the morphology of the Anw-body have a strong impact on AS and muscle organization. Anw is highly conserved in animals, and our results suggest that splicing regulation is the ancestral function of Anw. Remarkably, *anw* in all organisms studied harbors a conserved alternative exon allowing the coupling of AS and NMD for the regulation of this important splicing factor. Interestingly, one of Anw’s human homologs has been identified as a candidate gene for the muscle illness of Myasthenia gravis (MG) (Landouré et al. 2012), which suggests that AS mis-regulation could be a novel underlying mechanism for MG.

## Results

### Loss of Anw disrupts Myosin Heavy Chain production and flight muscle formation

We conducted a small-scale mutagenesis screen focusing on *Drosophila* genes encoding proteins with an annotated domain predictive of RNA functions (binding and/or enzymatic activities directed at RNAs) that have not been well characterized (e.g., Peng et al. 2023; Chen et al. 2024). CRISPR-Cas9 technique was employed to induce frame-shift mutations to the chosen genes. The *CG10948* gene was chosen because it has a single annotated RRM domain enriched in proteins with a function in RNA biology (Lasko 2000).

The initial round of mutagenesis led to the recovery of a single allele with an 8bp deletion leading to a premature stop codon, thus eliminating over 85% of the protein (Figure 1A). This allele of *CG10948* produces viable and fertile adults as either homozygous or heterozygous over a chromosomal deficiency (*df*) of the 69E6 region that contains *CG10948*. However, we noticed that both types of mutants (homozygous or hemizygous) display a wing posture phenotype in 100% of the adults. Majority of these flies have wings that, instead of extending out from the notum, drop to the sides of the body (the “droopy wing” phenotype, Figure 1B). In a minor fraction of the adults, particularly males, the wings are raised upward (Figure 1B). Based on this wing phenotype, we rename *CG10948* as *angel wing* (*anw*) and the initial allele *anw^1^*. In addition, *anw^1/1^* or *anw^1/df^*flies are flightless and unable to jump (Supplemental Video S1), suggesting that the responsible muscles are defective. We conducted two additional rounds of *anw* knock-out with Cas9 using two additional sgRNAs (Figure 1A). New *anw* frameshift allele produced identical “droopy wing” and flightless phenotypes as the original *anw^1^* allele. We therefore suggest that all of these frameshift mutations are likely nulls or severe loss-of-function alleles.

**Figure 1.**
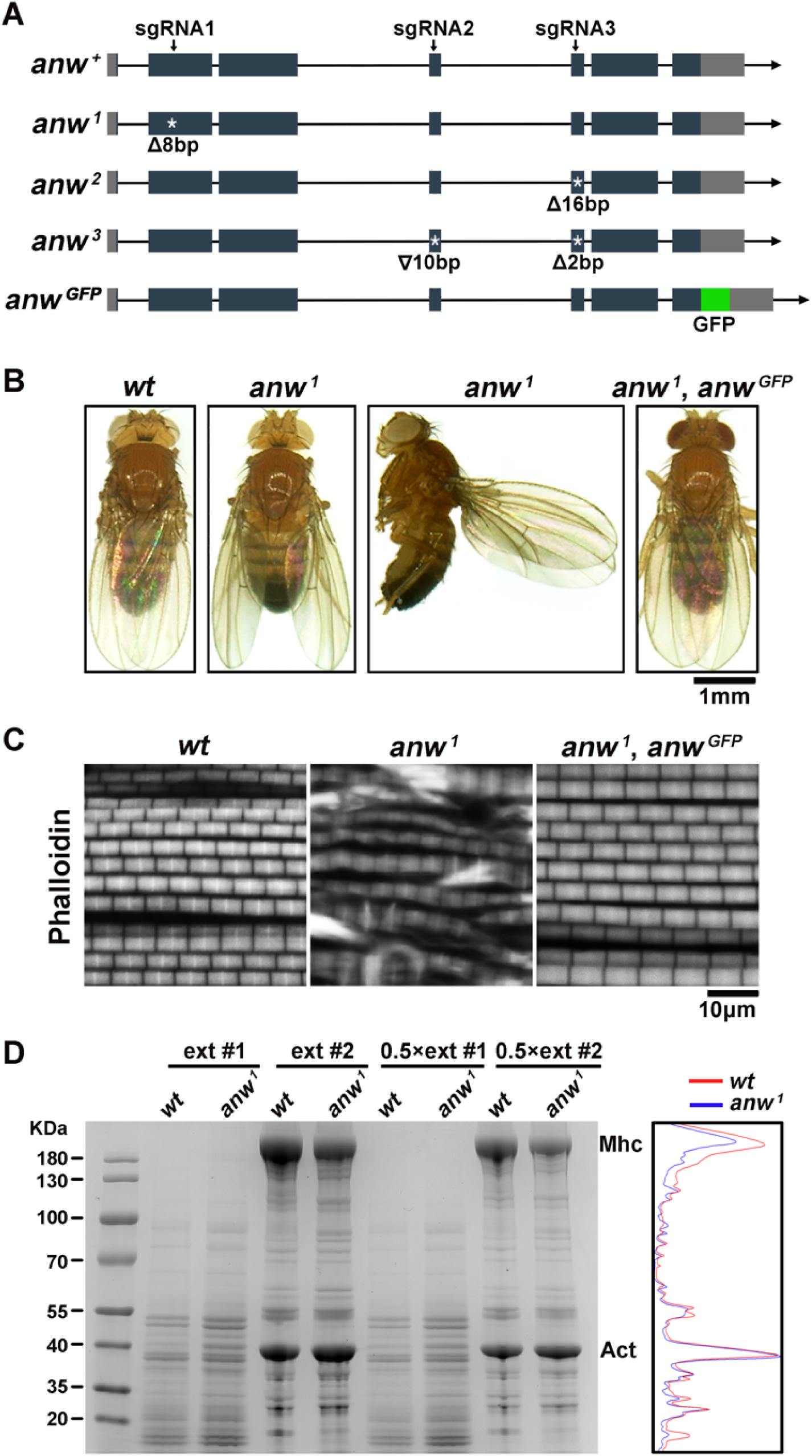
Wing posture and muscle defects in *anw*-mutant adults. **A. Alleles of *anw*.** In the diagram showing the genomic structure of the *anw* locus, rectangles represent exons, with coding exons shaded in black and UTRs in grey. Horizontal arrows indicate the direction of transcription. The positions of the three sgRNAs in Cas9-mediated mutagenesis are indicated by vertical arrows. White stars (*) mark the positions of the point mutations with the nature of the mutations described below. The green box represents the GFP tag in the rescue transgene. **B. Pictures of adult showing wing postures.** Genotypes are listed on top, and the red-eyed fly has the *anw^GFP^* rescuing transgene. **C. Phalloidin staining of adult muscle**. Genotypes are listed on top. **D. Coomassie-blue stained gel of differentially extracted muscle proteins**. Genotypes are listed on top. For details of the extracting method and the definition of extract #1 and #2 see Materials and Methods. The running positions of Actin and Mhc are labelled. Quantification is given to the right of the gel picture for the lanes where the extracts were loaded in half-strength (0.5x).

We observed massively disrupted muscle formation in phalloidin staining of actin filaments from the largest muscles in the thorax of adults (Figure 1C). Using a protocol based on previous approaches aimed at preferentially recovering muscle proteins (Cripps et al. 1994; see Materials and Methods), we observed a strong reduction of a protein species likely corresponding to Myosin Heavy Chain (Mhc), a major component of the muscle (Figure 1D). Therefore, *anw*-mutant flies suffer disruption of muscle formation likely caused by the reduction of Mhc protein levels, and possibly other muscle constituents.

We constructed a transgene that contains a 7.6kb fragment from the *anw* genomic region with its own upstream and downstream regulatory elements. A C-terminal GFP tag was added to Anw (Figure 1A). This transgene was able to rescue the wing and flightless phenotypes of all the *anw* alleles, and restore muscle formation to a pattern identical to that of a wildtype adult (Figure 1B, C). Therefore, we conclude that the mutant phenotypes are solely caused by the loss of Anw function. The GFP tag was later used to determine the cellular localization of Anw and its derivatives.

### Anw forms nuclear bodies

The Anw protein is homologous to the ENOX1 and ENOX2 proteins in human (Figure S1). The Enox1/2 proteins have been studied for over 30 years, and it was determined that they encode extra-cellular NADH oxidases in cultured human cells (Löw et al. 2012). This functional classification is difficult to reconcile with our mutational characterization suggesting that the *Drosophila* protein is important for muscle function, and specifically for the production of the Mhc protein. To shed light on this potential discrepancy, we studied the cellular localization of the *Drosophila* protein.

We first used adults that produce only GFP-tagged Anw proteins (full genotype *anw^1^ [anw^gfp^]/anw^1^ [anw^gfp^]*) so that GFP fluorescence could be used to localize Anw inside the cells. As shown in Figures 2 and S2, we observed primarily nuclear localization of Anw-GFP. This is true for all the tissues that we investigated including but not limited to the muscle, ovary and testis of adults, as well as the salivary gland of larvae. Remarkably, Anw-GFP is even present in the nucleus of the oocyte (Figure S2).

**Figure 2.**
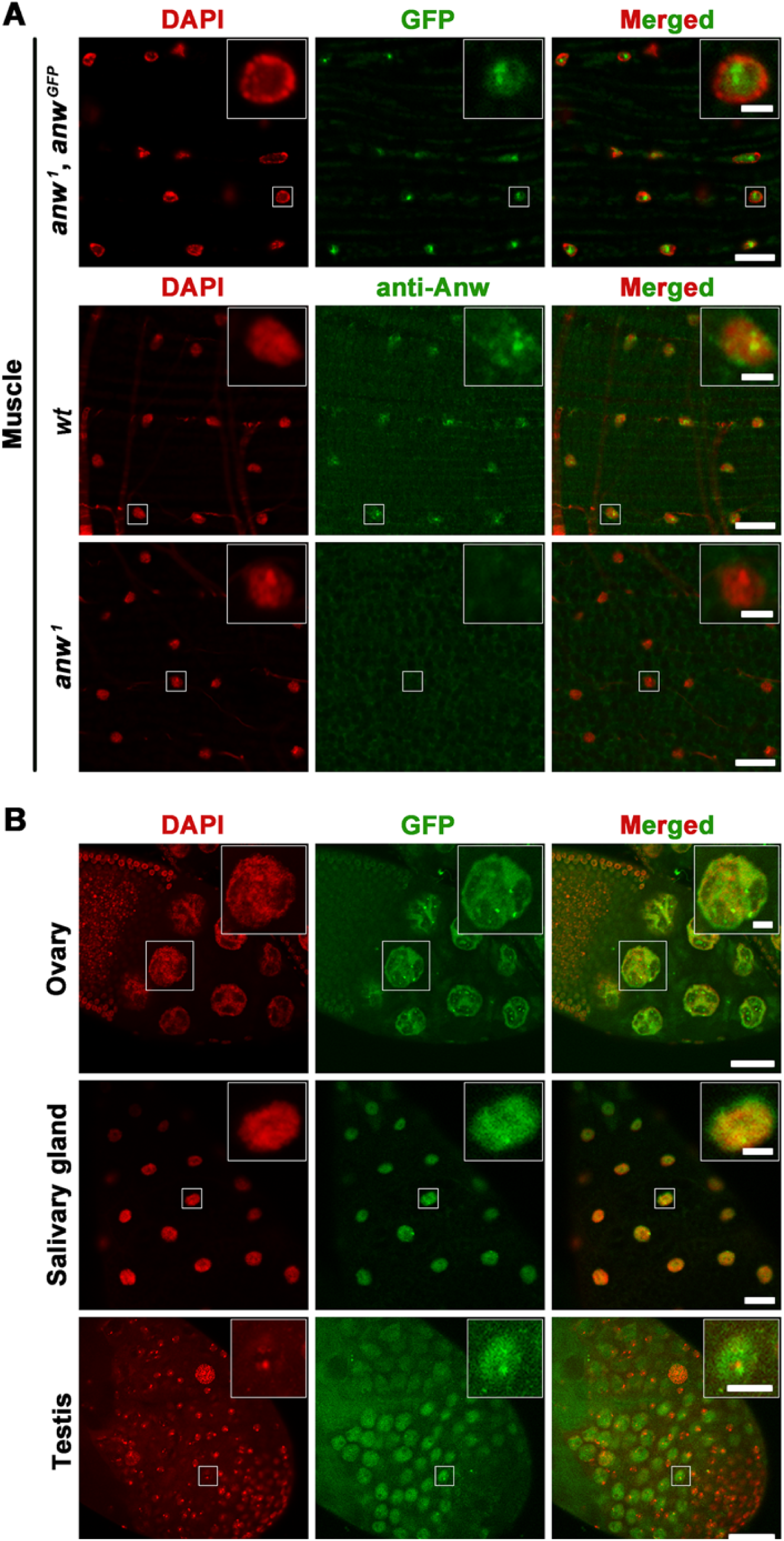
Anw localization in different tissues. **A. Anw localization in adult muscle**. The top panel (three images) shows GFP fluorescence (green) and DAPI (red) signals from an adult with the genotype shown to the left. One nucleus was chosen for which a magnified image is shown in the top right insert. In the lower panels (six images), anti-Anw staining of adult muscle with the genotypes listed to the left is shown. Scale bars are 2μm, and 10μm for the inserts. **B. Anw^GFP^ localization in other tissues.** The genotype was *anw^1^, anw^GFP^*. Scale bars are 4μm and 10μm for the inserts. Lower magnification images of Anw^GFP^ localization can also be found in Figure S2.

In addition, Anw-GFP forms nuclear speckles/foci in these tissues (Figure 2). In adult muscle cells, these foci are most prominent and seldom exceed two per nucleus. To rule out any potential artifact of using GFP-Anw as a substitute for Anw in these experiments, we raised antibodies against bacterially purified Anw antigens. These antibodies recognize foci in muscle cells of wildtype flies but fail to do so in *anw*-mutants (Figure 2A). Therefore, we conclude with confidence that *Drosophila* Anw protein is primarily a nuclear protein and forms intra-nuclear bodies. Anw’s specialized localization pattern and its possession of an RRM domain offer the first clues that Anw’s function might be related to RNA metabolism. We note that *Drosophila* muscle cells are enriched with foci formed by at least two other proteins important for pre-mRNA splicing, Aret/Bruno and Mbl (Oas et al. 2014; Spletter et al. 2015).

### Anw controls alternative splicing of introns enriched in neuromuscular functions

We performed whole transcriptome sequencing (RNA-seq) using two kinds of RNA samples, one recovered from whole adults of mixed sexes, the other from dissected thoraxes of adults of mixed sexes, which should be enriched with muscle tissues. As expected, we observed defects in *mhc* expression, particularly in the splicing of intron 9 of *mhc* (see below) in the mutant samples. We therefore conducted bioinformatic analyses profiling alternative splicing, and identified AS defects in multiple introns of different genes (see Supplemental Files S1 and S2). To manually verify some of the events defective in *anw*-mutants by RT-PCR, we implemented the following selection process. A group of 19 potential Anw-targeted regions from the “thorax” RNA-seq samples were selected with a fair representation of FDR ranks (Table S1). These 19 regions also passed visual inspection on the IGV genome browser for showing potential splicing defects in the mutants. 9 out of these 19 targets are also deemed splicing defective in the whole fly RNA-seq samples (Table S1). Using RT-PCR, we confirmed that 15 of the 19 selected show reproducible splicing pattern changes in the mutants *versus* the wildtype control. Importantly, splicing pattern changes in all of these 15 cases were rescued with the *anw^GFP^* transgene, lending strong support for the Anw playing an important role AS at these 15 regions. RT-PCR data for these 15 events are shown in Figures 3 and S3, with the genomic structure for each locus shown in Figure S4. Remarkably, 12 out of the 15 affected genes in *anw*-mutants have an annotated function in the nervous and/or muscle systems in the FlyBase. However, our results might have been biased as the thorax is expected to be enriched for neuromuscular tissues.

**Figure 3.**
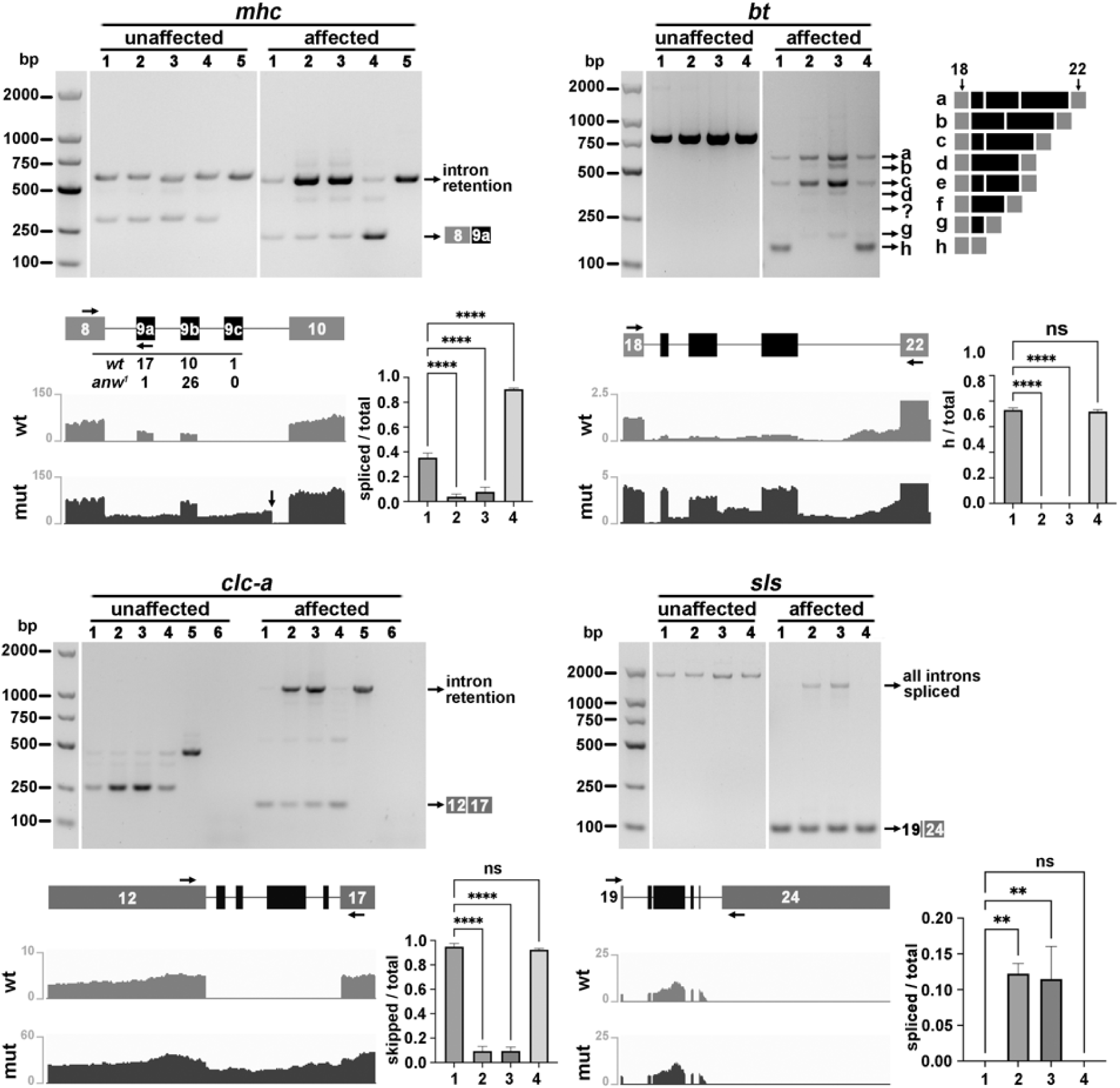
Alternative splicing events controlled by Anw. AS events in *mhc*, *bt*, *clc-a* and *sls* genes are shown. For the genomic structure of target genes see Figure S4. For each gene a gel picture is showed with results from two semi-quantitative RT-PCR reactions using total adult RNA. One primer pair spanning a region not controlled by Anw (“unaffected”) was used as a control. For their approximate positions see Figure S4. Another pair of primers span the “affected” region. The primer positions are indicated by horizontal arrows in the diagram of the exon/intron structure of the affected region, which is shown under the gel picture with exon labelling according to that in Figure S4. The products on the gel are either labelled with a description: “intron retention”, “all introns spliced” (for *mhc*, *clc-a* and *sls*), or with diagrams illustrating their structure. For *mhc*, the included exon (black) could be either type 9a or 9b or 9c. In the gel picture, a primer on 9a was used to demonstrate an enrichment of the 9a product in the *wt* but an enrichment of the retention product in the mutants. The distributions of the spliced products with different exons 9, recovered with another primer pairs, are tabulated under the exon/intron diagram of *mhc*. Under the exon/intron diagrams for all four genes are screen shots of the corresponding regions from an IGV browser loaded with RNA-seq reads. The vertical arrow in *mhc*’s IGV picture indicates the ectopic splice site utilized in the mutants. The genotypes for the lanes are as followed and indicated in parentheses: 1 (*wt*); 2 (*anw^1/1^*); 3 (*anw^3/df^*); 4 (*anw^1^, anw^GFP^*). For lane 5, genomic DNA from *wt* adults was used as the PCR template to indicate the running position of the un-spliced products. For lane 6 (in *clc-a* only), RT-PCR was performed with no reverse transcriptase added. Quantifications of a specific spliced product over total RT-PCR products are presented in the bar charts. ns: not significant; **: p<0.01; ****: p<0.0001. For additional Anw-controlled events see Figure S3.

Below we describe four Anw-controlled events in more detail as they will be investigated further. Others are described in Supplemental Result 1.

#### myosin heavy chain (mhc), exons 8-9-10

The *mhc* transcript is known to have complex AS patterns in different muscles and in different developmental stages (Nikonova et al. 2020). In particular, several clusters of exons undergo “mutually exclusive” AS (MXE) in which only one exon of a particular cluster is included in the mature RNA hence producing Mhc molecules with slightly different amino acid compositions at the site of the MXE exons (e.g., Bernstein and Milligan 1997; Standiford et al. 2001). Remarkably, many of these MXE exon clusters are conserved in Metazoan (Kollmar and Hatje 2014). In cluster 9, one of exons 9a, 9b and 9c is to be chosen, and the responsible mechanism likely involves long distance pairing of RNA elements (Yang et al. 2011). In *anw* mutants, we observed two classes of defective splicing. First, when we cloned and sequenced the products from RT-PCR using a pair of primers spanning the region between exon 8 and exon10, we uncovered a great over-representation of exon 9b (96% in *anw^1^* vs 36% in *wt*) at the expense of exon 9a in the mutants (Figure 3). Secondly, loss of *anw* led to elevated intron retention (RI) events between exons 8 and 10 (Figure 3). We noticed that the products with the appearance of RI actually contain splicing of an ectopic intron (arrow in the IGV browser picture). Interestingly this splicing event disrupts an important RNA element proposed to be essential for MXE at cluster 9 (Yang et al. 2011).

#### *bent (bt)*, exons 18-19-20-21-22

In the absence of Anw, the splicing event that results in skipping of all the intervening exons is largely missing, accompanied by an increase in the different classes of RT-PCR products representing the other AS events (Figure 3). Therefore, Anw appears to promote exon skipping for *bt* splicing.

#### chloride channel-a (clc-a), exons 12-13-14-15-16-17

Again, a drop of the event that skips all intervening exons was consistently observed in *anw* mutant samples. There appears to be a concomitant increase in RI in the mutants (Figure 3). To estimate any contribution from the contaminated genomic DNA in the RNA preparations, we included an RT-PCR test in which the reverse transcriptase was omitted. As shown in Figure 3, the PCR products corresponding to RI were essentially all templated from unspliced RNAs of *clc-a*. Therefore, the primary role of Anw for *clc-a* appears to be one that inhibits RI. Interestingly, there appears to be a concomitant increase in *clc-a* transcript level as evidenced from RT-PCR from the “unaffected” region and from IGV.

#### *sallimus (sls)* exons 19-20-21-22-23-24

Exons 20-21-22-23 are alternatively skipped, joining exons 19 and 24, in the wildtype. In the mutants, the level of skipping is reduced, giving rise to the splicing products in which all exons are precisely joined. These “all spliced” products have been verified by cloning and sequencing. Interestingly, this “mutant” splicing product is hardly detectable in total RNA from wildtype, implying that this product might be limited to specific tissues in wildtype animals. Our RT-PCR could not have detected the unspliced products due to their large size. The splicing defect in *sls* is not visually reflected in the IGV image for which we have no concrete explanation (Figure 3). However, multiple *anw*-mutant backgrounds consistently displayed this mutant pattern (see below), lending strong support that *sls* is one of Anw’s targets.

Although our molecular analyses were primarily focused on defects in exon-skipping in *anw* mutants, the clear presence of other types of splicing defect suggest that Anw does not control a unique class of AS events. Nevertheless, the precision by which Anw acts on its pre-mRNA target is remarkable and suggestive of targeting by RNA-protein interaction likely involving Anw.

### A mini-gene assay recapitulates Anw-dependent AS in cultured cells

To gain insights into how Anw regulates AS, we first attempted to recapitulate the Anw-controlled AS patterns in cultured *Drosophila* cells using transfected “mini-genes” producing the portion of pre-mRNA that Anw acts on in animals. The rationale is as follows. From RNA expression data available from the FlyBase, we surmise that Anw is not an abundant protein in *Drosophila* cells. Consistently, multiple antibodies raised against Anw were not able to detect the endogenous protein on Western blots, although the antibodies are effective in detecting Anw when overproduced (see below). We reasoned that by over-producing the pre-mRNA segments targeted by Anw, we might be able to achieve a situation in which the level of the endogenous Anw protein in the transfected cells is not sufficient to support normal splicing of RNA from the mini-gene, resulting in a splicing pattern similar if not identical to that in *anw* mutant animals. We further envisioned that if we could then provide additional Anw proteins ectopically in the above cells, we might be able to “restore” the AS pattern from the mini-gene to that of the wildtype flies, thus providing support for Anw’s direct involvement in regulating these AS events.

The overproduction in S2 cells of both proteins and pre-mRNA target segments was achieved using the Gal4-UAS system (schematics shown in Figure 4A). We first sub-cloned a full-length cDNA of *anw*, and the Anw-targeted genomic region separately, downstream of the UAS elements. A third plasmid carrying an *actin5C* promoter-driven *gal4* gene was used to produce Gal4, which activates transcription from the two UAS-bearing vectors. As a “no protein” control, we either transfected an “empty” UAS vector or one with an alternatively spliced *anw* cDNA clone that would encode a truncated protein missing the C-terminal 50% of Anw. Using Western blotting, we confirmed the overproduction of the full-length Anw protein (Figure 4B), but the presumed truncated protein was not detectable. Using the above setup, we succeeded in recapitulating the AS patterns for the Anw-targeted regions in *bt* and *clc-a* genes (Figure 4B). Remarkably, over-expression of the human Anw homolog Enox1 in *Drosophila* cells was sufficient to direct normal splicing (Figure 4B), providing the first line of evidence for functional conservation (see additional support below). Nevertheless, three other attempts (*aPKC, mhc, zasp67*) did not yield satisfactory results for reasons that evade our understanding.

**Figure 4.**
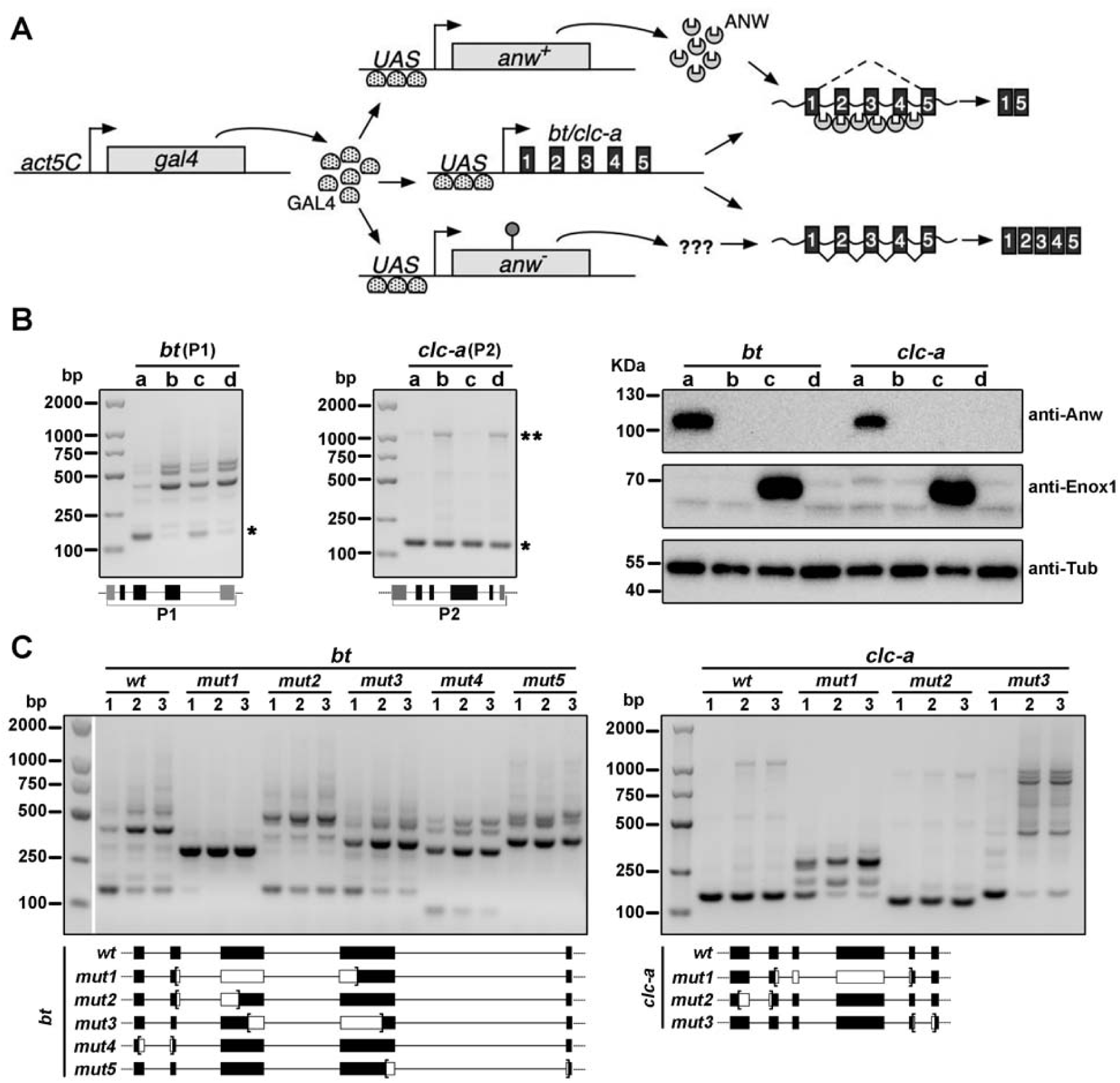
A mini-gene assay for the study of Anw-controlled splicing regulation. **A. Schematic representation of the mini-gene assay.** Straight horizontal lines represent DNA while wavery lines RNA. Boxes represent exons. Gal4 proteins produced from the *act5C* promoter act on *UAS* elements from different expression vectors. One produces full length Anw proteins, the other a presumed truncated protein due to the presence of a premature STOP codon (lollipop). These protein expressing vectors were individually transfected with an *UAS* driven mini-gene containing an alternatively spliced region of five exons. The mini-gene transcripts undergo two different modes of splicing (for illustration purpose only), depending on the presence or absence of the full length Anw protein. **B. Splicing regulation of *bt* and *clc-a* minigenes.** RT-PCR pictures are shown to the left using total RNA from cells transfected with the minigene reporters along with a plasmid expressing the full length Anw proteins (lane a), a prematurely terminated Anw isoform (lane b), the full length human Enox1 protein (lane c), and the empty expression vector (lane d). The primer pairs (P1 and P2) flanking the entire minigene were used to assay splicing. Differentially spliced products are marked with stars. See Figure 3 for their designations. A Western blot is shown to the right, to verify expression of the target proteins with antibodies listed to the right. **C. Mutational analyses of *bt* and *clc-a* minigenes**. RT-PCR gel pictures are shown using total RNA from cells transfected with the minigene reporters along with a plasmid expressing the full length Anw proteins (lane 1), a prematurely terminated Anw isoform (lane 2), and the empty expression vector (lane 3). In addition to the wild type minigene reporters, a series of internally deleted mutant minigenes were tested, with the diagram for the reporters shown under the gel picture in which the deleted regions are demarcated with brackets and shown in white.

To start deciphering the mechanism of action of Anw-dependent splicing, we first set out to identify whether any of the intronic regions of the minigenes is essential for Anw action, and generated a series of deletion constructs fusing neighboring exons. As shown in Figure 4C, deleting any single intron of the *bt* minigene affected AS of the minigene, although to different degrees, suggesting that there is not a single site of Anw action. A similar conclusion was reached for the *clc-a* minigene (Figure 4C). Interestingly, deleting the 3’ most *clc-a* intron led to the production of ectopic splicing products, but only in the absence of excess Anw proteins. Cloning and sequencing of these novel splicing products identified the usage of ectopic but genuine splicing sites in and around the largest exon. Although excess Anw seemed to suppress these ectopic splicing events, they were not detectable when a *clc-a* minigene was intact, either in the *anw* mutant flies or in transfected cells. It is therefore not clear to us how the regulation of these ectopic AS events relates to the function of Anw. As each deletion construct above is missing part of the exons flanking a particular intron (Figure 4C), a specific role of those exonic sequences in AS regulation can be ruled out. This leaves very few exonic sequences untested in our study.

In summary, we succeeded in establishing two different minigenes that when transformed into *Drosophila* cells reproduced an AS pattern that is controlled by the Anw protein and highly similar to that in wildtype animals. We have not succeeded however in identifying critical RNA element(s) within introns of the minigene that are necessary for Anw’s function as a splicing regulator.

Our working hypothesis is that Anw acts by binding to specific elements in pre-mRNA to achieve regulation of specific splicing events. This would be similar to other AS regulators with RNA binding capabilities (e.g., MBNL, Bruno/Arrest, ELAV, etc.). We set out to identify the binding sites of Anw by immunoprecipitation(IP)-based approaches. As multiple attempts of CLIP or RNA-IP followed by sequencing did not yield informative results using un-transfected S2 cells, we resorted to a minigene approach in which both the protein and its RNA targets were being overproduced. Even though we did not succeed in identifying any candidate motifs that Anw binds, we gained evidence from multiple approaches with the minigene system that supports a direct interaction of Anw with its RNA targets (see Supplemental Result 2 and Figure S5).

### Deletion mutations identify functionally important domains of Anw

To better understand Anw as an AS regulator, we conducted a structural and functional dissection aimed at identifying functionally important domains of Anw. A predicted structure shown in Figure S6 indicates that in addition to a well-conserved RRM domain, Anw possesses six helical structures and a poorly structured C-terminus. As the region N-terminal to RRM is poorly conserved (Figure S1), we started our deletion construction from the RRM domain. We first took advantage of the minigene assay established above, and identified potentially important domains for Anw function with nine mutant constructs (See Supplemental Result 3).

We next constructed a series of transgenes to introduce similar deletion mutations into flies for further characterizations (Figure 5A, for more details about the deleted residues see Figure S1). The designs were informed by the results from the minigene assay shown in Figure S6, and based on ease of cloning so not to involve multiple exons of the *anw* gene. In addition to the deletion mutations, we also included a point mutation of *anw* harboring several amino acid residues changes (Trp^454^Asp^455^ to Ser^454^His^455^, and Trp^468^ to Ser^468^), all residues invariant in Anw homologs. We fortuitously discovered that these combined changes severely disrupt Anw’s ability to direct AS of minigenes (Figure S6), and named this allele *anw^gTc^*. The amino acids changed in *anw^gTc^* also fall in the short helix #5, for which we did not construct a deletion mutation additionally. To facilitate localization studies of the mutant proteins, we tagged all mutants at the C-terminus with GFP, and introduced them individually into an *anw*-null background. A similarly tagged wildtype *anw* transgene was used as a substitution for the wildtype control. In subsequent analyses we focused on four aspects of Anw biology: the organismal effect on wing posture, the tissue-level effect on muscle formation, the molecular-level effect on splicing, and the cellular localization of the mutant proteins.

**Figure 5.**
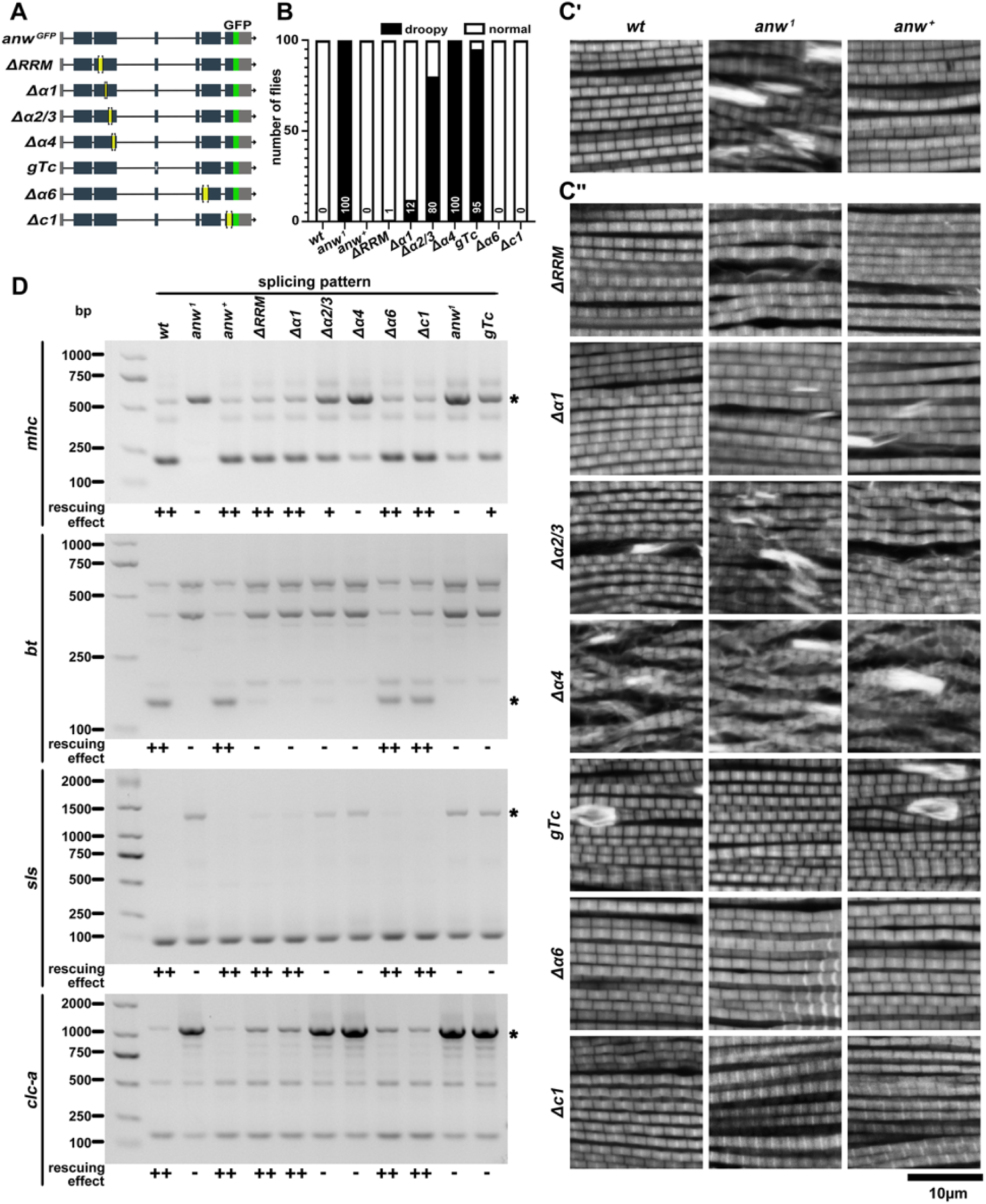
Domains of Anw important for muscle function and splicing regulation. **A. Transgenic constructs with mutated domains of Anw.** Designations for the constructs are given to the left. Boxes represent exons, and lines introns. Grey boxes represent UTRs, and black ones coding regions. The deleted regions are demarcated with brackets and labelled yellow. All transgenes are tagged with GFP (green box). The star (*) marks the approximate position of the *anw^gTc^* mutation. **B. Effect on wing posture**. One hundred adults of mixed sexes were scored for wing posture, and the number of flies displaying droopy wings is given for each genotype. All flies are *anw^1^*along with the rescuing transgenes listed under the chart. **C. Effect on muscle morphology**. Phalloidin staining of adult muscle is shown. In **C’**, a single image is shown for each genotype (given at the top) with *anw^+^* representing *anw^1^* mutant with a wildtype rescuing transgene. In **C’’**, three images are shown for each genotype (given to the left), showing varying degrees of abnormality. **D. Effect on splicing**. The splicing patterns from *mhc*, *bt*, *sls* and *clc-*a genes are used to assay the effects of *anw* mutations. For each gene, a gel picture was shown for RT-PCR analyses similarly to ones shown in Figures 3 and S3 with differentially spliced products marked with stars. The “rescuing effect” is estimated from the RT-PCR data. Genotypes are listed on top.

#### The *anw^ΔRRM^* mutation

At the organism level, the droopy wing phenotype is greatly rescued even when the RRM domain is deleted (Figure 5B). Consistently, the pattern of major muscle in phalloidin staining experiments is more similar to the wildtype than to that from of the null mutations (Figure 5C), although defects of varying degrees remain. At the cellular level, deleting the RRM domain does not alter the nuclear localization ability of the protein (Figure 6A and B). We stained muscle from these flies with anti-Anw, and observed co-localization of the antibody and GFP signals (Figure 6A). However, nuclear GFP foci of the mutant protein appear brighter and more compact than wildtype Anw^GFP^, which is most prominent in testes and ovaries (Figure S7). In addition, there appears to be less GFP signals outside of the focus, indicating that the mutation might have increased the protein’s ability to congregate thus reducing the level of the “free” protein. These changes in protein localization brought about by the *anw^ΔRRM^* mutation are most evident in muscle and primary spermatocytes in the testis (Figures 6 and S7). However, without an accurate estimation of the mutant *versus* the wildtype expression levels, we cannot rule out that these pattern changes to some degrees reflect the changes in protein levels. At the molecular level, we chose four Anw-regulated splicing targets (*clc-a*, *bt*, *sls*, and *mhc*) to assay the effects of the *anw* deletion mutations. As shown in Figure 5C, loss of RRM partially abolishes splicing regulation of some but not all of the chosen targets. These results suggest that the RRM domain is important but not absolutely required for splicing regulation, which is consistent with the partial rescue of the organismal phenotypes.

**Figure 6.**
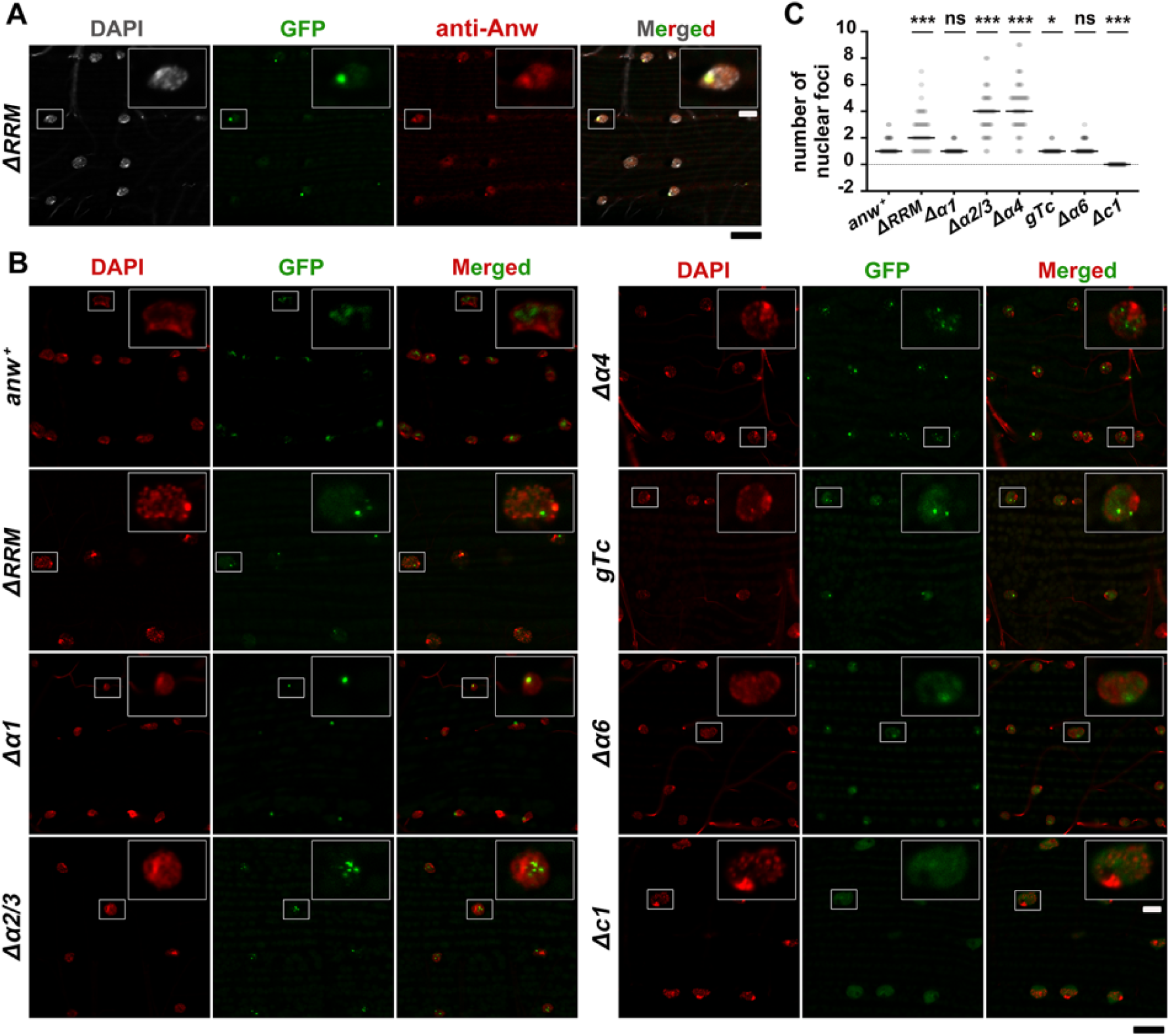
Domains of Anw important for nuclear body formation in the muscle. **A. Assessing Anw localization with GFP florescence and antibody staining**. Adult muscle expressing GFP tagged Anw^ΔRRM^ protein was stained with an anti-Anw antibody, showing DAPI signal in white, GFP fluorescence in green, and antibody signal in red. A single nucleus outlined in white was chosen for presenting a magnified image shown in the top right insert. **B**. **Localization of different Anw mutant proteins in muscle cells**. Using GFP fluorescence, the localization of Anw mutant proteins was studied. Genotypes are listed to the left. **C. Quantification of the number of nuclear foci in different mutants**. The number of Anw foci in muscle nuclei are counted in adults expressing different Anw mutant proteins. The median numbers (highlighted in black) were compared with that of the wildtype by a Mann-Whitney test. ns: not significant; *: p<0.05; ***: p<0.001. Scale bars are 2μm and 10μm for images in inserts.

#### The *anw^Δα1^* mutation

About 90% of the adults with this mutation no longer have droopy wings (Figure 5B). Flight muscle patterns are similar to those displayed by adults with the *anw^ΔRRM^* mutation (Figure 5C). The protein localization patterns are similar too (Figures 6B and S7), so are the degrees of rescue on splicing (Figure 5D), for the two mutations. The two affected domains juxtapose each other on the Anw protein (Figure S6B). The remarkably similar degrees of phenotype lead us to suggest that they are two parts of a larger functional domain.

#### The *anw^Δα2/3^* mutation

About 80% of the adults with this mutation display droopy wings (Figure 5B). Muscle patterns are more severely disrupted than those with the first two mutations (Figure 5C), although less so when compared to the null mutants. Consistently, disruption to splicing appears more severe than the first two mutants (Figure 5D). Interestingly, localization of this mutant protein deviates from the normal pattern in that there is an increase of Anw foci per muscle nuclei from a median of 1 focus in wildtype to 4 in the mutant (Figure 6C). Unlike the first two, the *anw^Δα2/3^* mutation results in a specific reduction or increased diffusion of the protein in salivary gland nuclei from third instar larvae (Figure S7). Since the regulatory elements for proper gene expression remain the same for all transgenes, and each was inserted at the same chromosomal location of 86F on chromosome 3R, the differences in protein level must reflect different behavior of the mutant proteins.

#### The *anw^Δα4^* mutation

This mutation resulted in the strongest mutant phenotypes (least degree of rescue) amongst all the deletion mutations that we tested (Figure 5). Interestingly, the mutant protein assumes a localization pattern similar to that from the *anw^Δα2/3^* mutants, in that Anw appears in multiple foci in multiple tissues (a median of 4 foci in muscle nuclei, Figure 6C), except the salivary gland where protein level is similarly reduced or diffused as in *anw^Δα2/3^* (Figure S7). The two affected domains juxtapose each other on the Anw protein. Based on the similar phenotypes displayed by their disruptions, they might represent two parts of a larger domain, with a main function of preventing the formation of excessive number of Anw focus.

#### The *anw^gTc^* mutation

This mutation is the most subtle in that it only changes three residues but is one of the more severe *anw* alleles. Over 90% of the adults have droopy wings (Figure 5B). However, muscle disruption appears weaker than *anw^Δα2/3^* (Figure 5C). The localization pattern of Anw^gTc^ is similar to that of the wildtype protein (Figures 6B and S7) in most tissues, except in nurse cells of the ovary where its level is greatly reduced, and in salivary gland cells where it loses the ability to form foci (Figure S7). Contrary to its near normal nuclear localization, this mutant confers one of the weakest degrees of rescue on splicing defects (Figure 5D).

#### The *anw^Δα6^* mutation

In this weak mutant, despite defective muscle patterning still being evident (Figure 5C), all adults examined have normal wing posture and splicing regulations are largely restored (Figure 5D). Despite the mild organismal defects conferred by the mutation, Anw^Δα6^ protein has a disrupted localization pattern (Figures 6B and S7), in that nuclear foci become less prominent and sometimes absent, particularly in polyploid nuclei such as those in salivary glands and the nurse cells (Figure S7). Therefore, although partially dispensable for the maintenance of muscle structure and AS regulation, helix 6 is involved in efficient focus-formation of Anw, which suggests that the formation of nuclear foci *per se* is not absolutely required for Anw functions.

#### The *anw^Δc1^* mutation

The prior conclusion that nuclear foci formation is not an absolute requirement for Anw functions is further supported by data collected from the *anw^Δc1^* mutation, which is essentially the identical to *anw^Δα6^* in its ability to rescue *anw*-mutant phenotypes (Figure 5). Remarkably, losing the 60-aa C-terminal domain largely abolishes Anw’s ability to form foci but not localize to the nucleus. Again, we suggest that helix 6 and the C1 domain with their relative proximity constitute a single functional domain. This proposition is consistent with results from cultured cells (Figure S6), where expressing the Anw with a C2 deletion, encompassing both the helix 6 and C1 regions, does not affect normal splicing of both the *bt* and *clc-a* mini-genes.

In summary, our mutational studies uncovered functional domains in Anw, which we tentatively grouped into four regions. The first comprises the RRM plus helix one, which are required for maintaining a less condensed appearance of the Anw nuclear foci. Disruption of this region has an intermediate effect on muscle development or splicing regulation. The second region comprises helixes 2 to 4, which are required for maintaining the appearance and number of the Anw nuclear foci. Disruption of this region has the strongest negative effect on the function of Anw. The third region is composed of the short helix 5, which plays a minor role in controlling the behavior of the protein inside the nucleus. Remarkably, this region when disrupted has a very strong effect on splicing. The fourth region is composed of the C terminal third of the protein, whose primary function seems to be allowing congregation of the protein inside the nucleus. Interestingly, disruption of this large region affects splicing sparingly.

However, there are clear inconsistencies revealed by our genetic analyses. First of all, mutant proteins can assume different distribution modes in different tissues. We currently do not understand the underlying mechanism but wish to suggest that it may involve other proteins that facilitate Anw body formation as well as their distribution in different cells. Secondly, disruption of muscle patterns does not necessarily correlate with the severity of splicing disruption. There is precedent for inconsistency in muscle morphology and muscle function. For examples, normal muscle morphology but not function was restored in other studies when Mhc isoforms were inappropriately expressed (Wells et al. 1996; Swank et al. 2001). A more consistent picture might emerge when we investigate more Anw targets and/or dissect the effects in a tissue-specific manner.

### Anw is a class of AS regulators in animals with dynamic evolution

Although there is only one *anw* homologous gene in *Drosophila*, there are two in human, *Enox1* and *Enox2*, which share similar degrees of homology to Anw (Figure S1). We are interested in the evolutionary trajectory of the Anw class of proteins and conducted BLAST searches for Anw homologous proteins in the databases. We discovered that Anw has a specific and ancient presence in animals (Figure 7A), with the sponge (Porifera) being the earliest phylum of its presence. Interestingly, the *anw* gene experienced multiple losses during evolution, as evidenced by its absence in both Nematoda, which includes the model worm *C. elegans*, and Platyhelminthes, which includes the planarians. A recent loss might have happened in the ancestor of Urochordata since *anw* is absent in Ciona. Moreover, in Cyclostomata, a single *anw* is present in the lamprey *Petromyzon marinus*, but absent in its sister species, the Hagfish. We cannot rule out the possibility that the absence of *anw* in these less studied organisms is just due to gaps in genomic sequencing.

**Figure 7.**
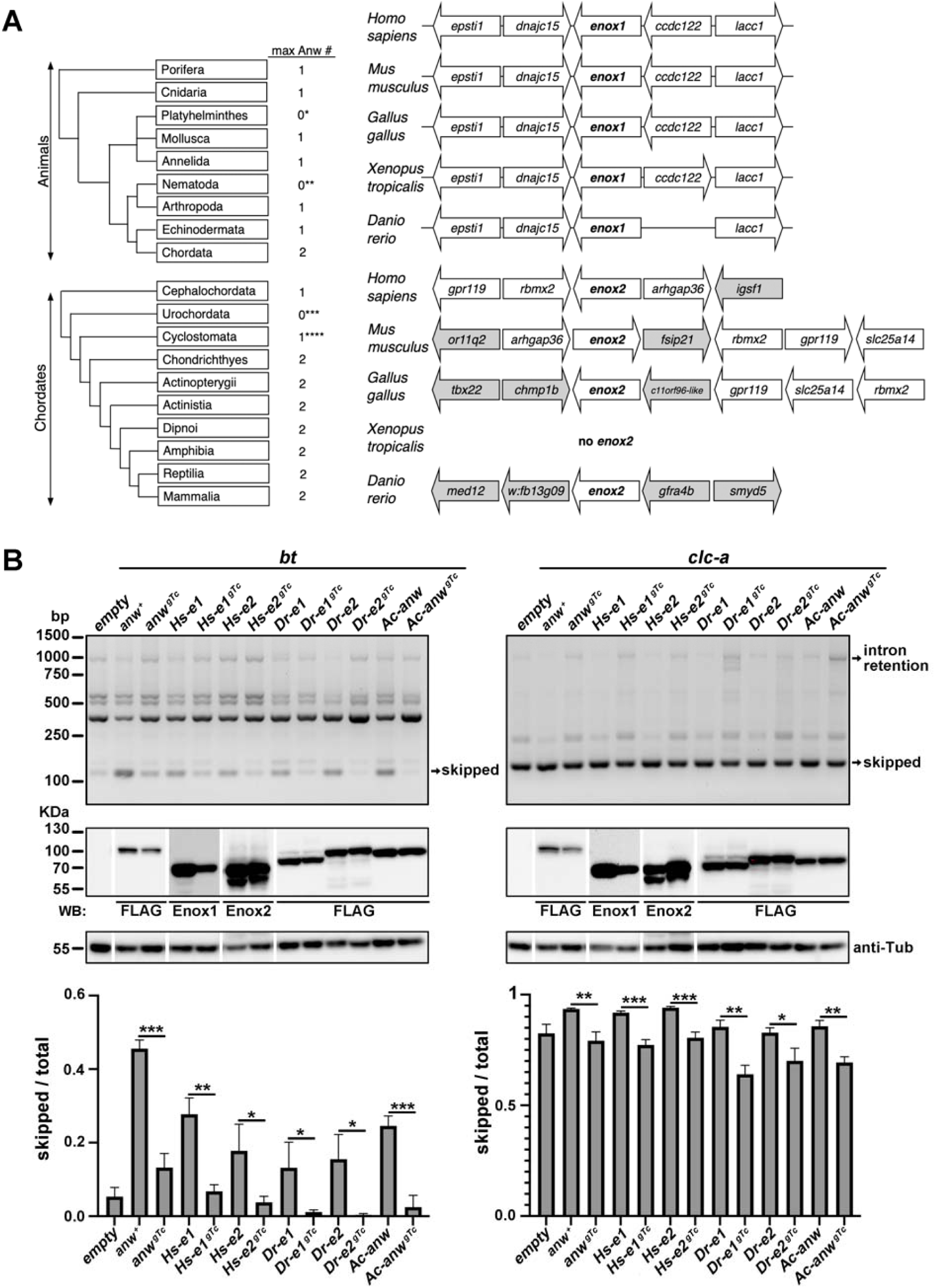
Anw homologous proteins are functional in *Drosophila* cells. **A. Gene gains and losses of *anw* during animal evolution**. At the left are phylogeny trees given for animals and chordates along with the maximal number of Anw homologous genes present. *: phylum includes the planarians; **: phylum includes *C. elegans*; ***: phylum includes Ciona. ****: one *anw* gene is present in lamprey but none in the Hagfish. At the right is synteny analysis of human *enox1* and *enox2*. Genes are represented as block arrows with the arrow indicating the direction of transcription. Genes in white display syntenic relationship with either *enox1* or *enox2* in the five vertebrate species chosen. Genes in grey are out of synteny. **B. Anw-homologous proteins direct splicing in *Drosophila* cells**. The effect of expressing Anw homologous proteins in S2 cells was tested on splicing from the *bt* and *clc-a* minigenes. For each minigene, the top panel shows a gel picture of RT-PCR analysis for splicing. The middle panel shows Western blotting verification of the expression of the target proteins with the antibodies used indicated below the pictures. As the same membrane was probed with multiple antibodies, the molecular markers were used for alignment in picture assembly. The bottom bar chart shows quantification of splicing products from three replicate experiments. Hs: *Homo sapiens*; Dr: *Danio rerio*; Ac: *Aplysia californica*; Empty: empty expression vector; e1: *enox1*; e2: *enox2*. gTc: the equivalent mutation as the *Drosophila anw^gTc^* mutation. ns: not significant; *: p<0.05; **: p<0.01; ***: p<0.001.

The *anw* gene experienced at least one recent gain in Chordata, giving rise to the human *Enox1* and *Enox2* genes. A simple synteny analysis suggests that *Enox2* is the more ancient copy of the two, based on fewer changes of genes immediately adjacent to *Enox*1 than to *Enox2* (Figure 7A). However, evolutionary age analysis using the recently developed GenTree program (Shao et al. 2019) did not yield a large age difference between the two genes. Nevertheless, it confirms a dynamic evolutionary trajectory for the *anw*-class genes. Expression data available in public databases suggest that the two genes have different expression patterns. *Enox2* is widely expressed while *Enox1*’s expression is more restricted. For example, Mouse ENCODE transcriptome data seem to suggest that *Enox1* is enriched in the nervous system (https://www.ncbi.nlm.nih.gov/gene/239188), while *Enox2* has a much broader expression (https://www.ncbi.nlm.nih.gov/gene/209224). This would be consistent with that *Enox2*’s ancestral origin that we suggested earlier from the analysis of synteny, and with the broad expression of *anw* in *Drosophila*.

The observation that Anw is widely present in animals and has tissue specific paralogs in higher organisms begs the question of whether its ancestral function is in splicing regulation. Encouraged by that cross-phylum rescue experiments can yield positive outcomes (Vicente et al. 2007; Vicente-Crespo et al. 2008) and by our results with Enox1 directing direct splicing of *Drosophila* minigenes (Figures 4B and S6C), we took advantage of the mini-gene system to preliminarily test this hypothesis. We expressed human Enox2 in *Drosophila* cells, and showed that it is also able to direct splicing of *Drosophila* minigenes (Figure 7B). As negative controls, we generated single mutations to the three conserved residues in Anw homologs (the *anw^gTc^* mutation, Figures S1) in the human Enox1 and Enox2 proteins. As expected, neither of the mutants was able to direct splicing, supporting our hypothesis that the Anw class of proteins are splicing regulators. We were encouraged with these results and tested additional Anw homologs from Aplysia, zebra fish (both homologs), and obtained essentially identical results, each case supported by the same *anw^gTc^* mutation as a negative control (Figure 7B). These results lend strong support that the ancestral function of Anw proteins is likely in splicing regulation.

### A cross-phyla ultra-conserved DNA element is a potential autoregulator of Anw

The residues mutated in the strong *anw^gTc^* mutation lie in a small exon (107bp) of the *anw* gene that displays remarkable features of evolutionary conservation. As shown in Figure 8, not only is this small exon present in all the *anw* homologous genes that we investigated, its length and DNA sequence are also exceptionally conserved through animal evolution.

**Figure 8.**
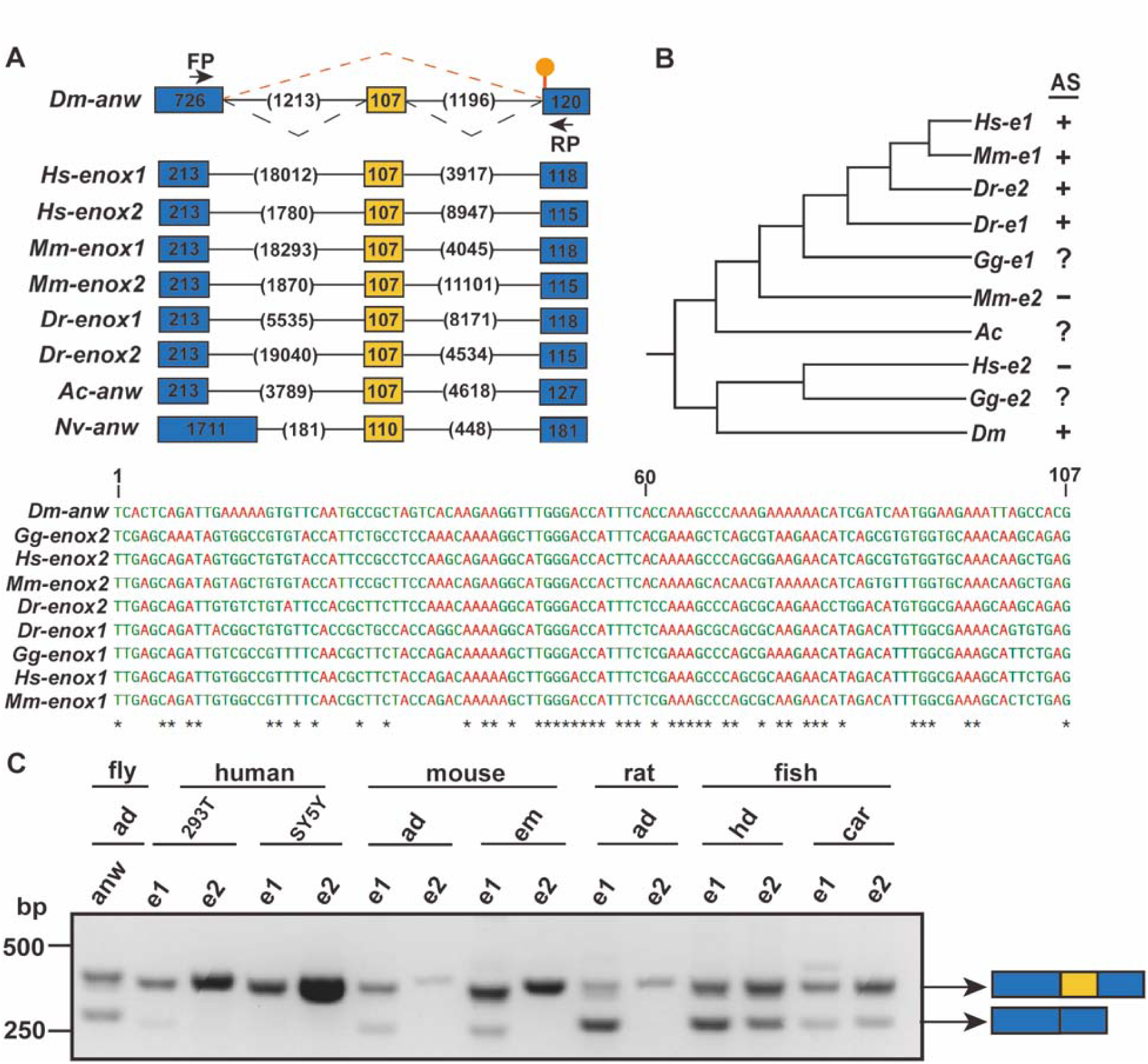
An ultra-conserved element as an alternative exon for potential *anw* regulation. **A. A conserved exon in *anw* homologous genes**. The exon/intron structure of the region with the small exon labelled yellow, and the immediate up- and down-stream exons in blue. Their sizes in bp, and the sizes of the intervening introns are shown. The position of the pre-mature STOP codon in mRNA species without the exon is marked with a lollipop. Dm: *Drosophila melanogaster*; Hs: *Homo sapiens*; Mm: *Mus musculus*; Dr: *Danio rerio*; Ac: *Aplysia californica*; Nv: *Nematostella vectensis*. **B. The small exon is highly conserved in DNA sequence**. A comparison of the exons from *Drosophila* and other sequences is shown as a tree. Gg: *Gallus gallus*. The sequence alignment used to construct the tree is shown below. PCR results from **C** were used to infer whether the exon is alternatively spliced (AS). **C. Alternative splicing of the small exon**. RT-PCR with primers in the flanking exons (FP and RP in **A**) were performed on total RNA isolated from human cell lines (293T and SY5Y), and different tissues from three other species: adults (ad) from fly, mouse and rat; embryos (em) from mouse; head (hd) and carcass (car) from zebra fish. The “skipped” (lower bands) and “retained” (upper bands) products are labelled.

This exon is similar to a rare class of DNA elements termed “ultra conserved element” (UCE), first defined in mammals as a stretch of about 200bp of DNA having over 99% identity between mouse and human (Bejerano et al. 2004). This 107bp exon is 99% identical between mouse and human *enox1* (95% for *enox2*), 70-80% identical between fish, chicken and human. In *Drosophila*, similar UCEs (100% identity within *Drosophila*) have been described (Kern et al. 2015). Remarkably, one of the *Drosophila* UCEs in the coding region include this 107bp exon in *anw*.

UCEs serve important function as splicing regulators. It was noticed over 15 years ago that SR splicing factors in mammals are encoded by genes with alternative exons consisting of ultra or highly conserved sequences (Lareau et al. 2007; Ni et al. 2007). The inclusion of these exons inevitably creates a pre-mature stop codon leading to the non-sense mediated degradation (NMD) of the transcript. Such a mode of regulation coupling AS and NMD seems to be highly conserved (Lareau and Brenner 2015). Interestingly, the sequence conservation of the “poison exon” in a particular SR gene family usually does not extend beyond its phylum, unlike the small exon in *anw* genes.

We propose that this cross-phyla UCE fulfills a similar regulatory function mediated by the combined action of AS and NMD. Hansen et al. (2009) showed that *anw* is one of the *Drosophila* genes subject to NMD regulation, via the inclusion/exclusion of this ultra-conserved exon. We performed RT-PCR tests on samples from a variety of different organisms, and obtained evidence supporting AS of this exon in *enox1* of all tested organisms and *enox2* in zebrafish (Figure 8). This seems to contradict the ancestral origin of Enox2 as we suggested earlier, again reflecting the complexity of Anw evolution. We notice a deviation in AS-NMD regulation of *anw* from the theme observed for the SR genes. In SR genes, the alternative exon serves to create a premature stop codon when included, whereas the inclusion of 107bp exon in *anw* restores the reading frame allowing the production of a full length Anw protein. This potential mode of Anw regulation suggests that the level of Anw protein is tightly regulated *in vivo*. Consistently with this hypothesis, when we overproduced Anw in the muscle of otherwise wildtype adults by an *mhc*-Gal4 driven expression, all adults developed droopy wings (n>100). The underlying defects in this Anw overexpression background await future investigations.

## Discussion

Here we characterized the founding member of a class of conserved alternative splicing regulators, the Anw protein. Anw shares some of the known features of splicing factors, such as controlling a specific set of AS events, and having the propensity to interact with RNA targets as well as the ability to multimerize in the nucleus. Importantly, Anw and possibly all of its homologs employ the coupling of AS and NMD to control its expression level. Except for an annotated RRM domain, Anw and its homologs share no domain similarity with other classes of known AS regulators, including the SR proteins or the hnRNP proteins. Therefore, further elucidation of how Anw acts is likely to reveal new principles governing splicing regulation.

### How Anw regulates splicing: protein-RNA interaction

It is clear that Anw only controls a small subset of the AS program in *Drosophila* tissues. The most parsimonious model is that this precision is determined by Anw’s recognition of its target sequence/structure on pre-mRNA, similar to other known RBP splicing regulators, such as MBNL, Bruno/CELF, ELAV/Hu, Pasilla/Nova. Even though we presented evidence supporting Anw’s ability to interact with its target RNA molecules, we failed to identify a candidate consensus for this recognition. Moreover, considering that the RRM domain is the prime candidate for Anw’s RNA binding ability, we were surprised to discover that an RRM-less Anw retains partial ability to direct splicing *in vivo*. These results suggest a few possible scenarios. First, the primary role of RRM in Anw is not to interact with RNA, but to interact with proteins. Such precedence exists. For example, both empirical and computational analyses suggest the existence of at least six cases where an RRM is primarily involved in protein-protein interactions (Maris et al. 2005; Shazman and Mandel-Gutfreund 2008). In this case, Anw’s RNA binding ability might be brought about by other domain(s) of the protein. A role of RRM in protein-protein interaction is also supported by our results from studying the localization of the mutant proteins (see discussion below). Secondly, RRM along with another domain jointly control Anw-RNA interaction so that losing just the RRM is not sufficient to completely abolish binding. Such precedence also exists. For example, the Intrinsically disordered RGG/RG domains often juxtaposing RRMs have RNA binding activity, and together with RRM establishes high specificity binding (Ozdilek et al. 2017). Structural prediction suggests that almost the entire region C terminal to the RRM in human Anw homologs constitutes a domain architecture homologous to the “structural maintenance of chromosome segregation protein SMC, common bacterial type (SMC_prok_B)” in NCBI database (TIGR02168). SMC proteins control chromosome segregation by binding DNA/chromatin. However, the appearance of such a domain in a splicing regulator has no precedence. Thirdly, Anw does not interact with RNA. It is recruited to its targets by interactions with another RNA-binding protein(s). It is worth noting that RRM in Anw belongs to a special class in which the only member is found in Anw and its homologs. Further characterization of this class of RRM, and Anw’s interacting partners would help distinguish the above hypotheses.

### How Anw regulates splicing: nuclear body formation

Anw displays a property shared by other splicing regulators: the ability to form nuclear foci/speckles. Examples of such splicing regulators include the Bruno/Arrest (Oas et al. 2014; Spletter et al. 2015), ELAV (Yannoni and White 1997), MBNL (Vicente-Crespo et al. 2008; Oas et al. 2014), and Caper (Titus et al. 2021) proteins in *Drosophila*. A simple notion emerges that the foci/speckles are places where splicing factors/activities enrich (Barutcu et al. 2022; Bhat et al. 2024; Wu et al. 2024). We showed by domain deletion that the C terminal one-third of Anw, largely disorder in structure, is essential for focus-formation. Paradoxically, C terminally deleted *anw* mutations display the weakest loss-of-function phenotype in muscle, and support splicing *in vivo* and in cultured cells to a large extent. These results suggest that nuclear focus formation is not absolutely required for Anw’s function. However, we cannot rule out that smaller foci beyond the resolution of our microscopy analyses are still being formed by C-terminally deleted Anw proteins. How the C-terminus of Anw functions might become clearer once we have discussed deletion analyses of the rest of the protein.

Structural prediction suggests that the region adjacent to the RRM domain is made of several alpha helixes (flybase.org). Individual disruption of those helixes all leads to defects in nuclear body formation and splicing regulation, although to varying degrees. In particular, deletions within the region including the RRM domain result in Anw foci becoming more abundant and with a more condensed appearance. Remarkably, these mutations have the strongest effect on Anw’s function with the *anw^Δα4^* mutation behaving essentially as a null. These results lead us to speculate that these mutant proteins are trapped in a non-functional state. It has been previously proposed that nuclear bodies of splicing factors, such as Bruno/Arrest and MBNL, serve as their storage granule (Oas et al. 2014). Our proposition above appears consistent this model: aberrant storage formation might result in trapping of the components making a complete loss of function possible, whereas dismantling the storage (caused by the C-terminal deletions for example) would only weaken the function as active components are still available even in lower quantities. As suggested earlier, RRM in Anw is primarily a protein interacting domain. Its loss affects morphology of Anw bodies, which would be consistent with RRM having a role in mediating Anw multimerization, a role also assigned to RRMs in other RBPs (e.g. Toba and White 2008; Zacco et al. 2018).

In summary, Anw shares an interesting nuclear localization property with other splicing regulators. Nevertheless, how nuclear body formation relates to its molecular function in splicing regulation awaits further investigations.

### Is Anw/Enox1/Enox2 a case of protein moonlighting or mistaken identity

Our conclusion that the Anw protein functions as an intra-nuclear factor important for RNA splicing contradicts the annotated functions of Enox1 and Enox2 in human. Both proteins are believed to be outer cell surface-attached NADH oxidases with at least two activities demonstrated *in vitro*: a hydroquinone (NADH) oxidase activity and a protein disulfide-thiol interchange activity (https://www.ncbi.nlm.nih.gov/gene/55068 and https://www.ncbi.nlm.nih.gov/gene/10495). Therefore, we might have on hand a case of protein moonlighting in which a single polypeptide fulfills more than one functions that are unrelated, and those distinct activities happen at different cellular spaces (reviewed in Jeffery 2018; Singh and Bhalla 2020). However, a careful survey of the literature raised several concerns for this hypothesis.

The determination of the above NOX enzymes being extra-cellular was originated from antigen mapping of a monoclonal antibody raised against a “cell surface glycoprotein” enriched in various tumor cell lines (Chang and Pastan 1994). However, new antibodies raised against the bacterially purified antigen protein (a fragment of Enox2) failed to recognize the cell surface antigen. Secondly, an outer cell surface associated NADH oxidase was shown sensitive to drug inhibition without identifying the molecular nature of this enzyme (Morré 1995). In a later study, a cell surface associated NADH oxidase activity was purified, and one or more of the components reacts with antibodies raised against either tNOX (Enox2) or cNOX (Enox1) antigens (Scarlett et al. 2005). Unfortunately, opportunities were repeatedly missed to unambitiously attribute the ecto-NOX activities to Enox2 as multiple studies reporting efficient RNAi knocking down of its expression did not touch upon the important topic of whether the cell surface associated NADH oxidase activity is concomitantly reduced (e.g., Islam et al. 2024).

On first appearance, the Enox1/Enox2 case fits well with many cases of protein moonlighting in which a metabolic enzyme displays unexpected additional functions (Chen et al. 2021). However, in those cases, the enzymes have a clearly identifiable functional domain(s) associated with the enzymatic activity, which is not the case for either Enox1 or Enox2. Mutational studies of Enox2 were nevertheless carried out but limited to using bacterially purified proteins (Chueh et al. 2002). Although residues critical for some of the enzymatic activities *in vitro* were identified in the study, their importance *in vivo* has not been investigated.

Lastly, prominent localization of Enox1 or Enox2 to human cell surface lacks experimental proof (www.proteinatlas.org). Results from our Anw localization study suggest that *Drosophila* Anw is primarily if not exclusively nuclear. To demonstrate that a small amount of Anw proteins is targeted to the cell surface might require an Anw mutation blocking such transport, which would expectedly result in the accumulation of Anw in the cytoplasm. We have disrupted eight domains conserved between the *Drosophila* and human proteins. All of them results in some degrees of loss of Anw function, but none alters Anw’s nuclear localization, consistent with that the ancestral localization pattern of Anw class of proteins is primarily in the nucleus.

Therefore, whether the human Enox1 and Enox2 are moonlighting as extra-cellular oxidases requires substantially more vigorous investigations. In fact, concerns about some of the claims on Enox1/2 functions and localization have been previously raised (Löw et al. 2012). Nevertheless, our results demonstrating that the Aplysia, fish and human Anw homologs are functional in directing splicing in *Drosophila* cells favors the notion that at least the ancestral function of this class of protein is in splicing regulation.

### How the loss of Anw could have caused familial myasthenia gravis

AS are most abundant in mammalian nervous and muscular systems, consistent with many neuromuscular diseases are associated with underlying defects in RNA splicing. Myasthenia gravis (MG) is largely considered as an autoimmune disease of the neuromuscular junction (reviewed in Koneczny and Herbst 2019; Vilquin et al. 2019; Kaminski et al. 2024), and defects in AS have not been widely suspected to cause MG. In 1994, cases from a large Italian family suffering from late-onset MG were reported (Bergoffen et al. 1994). Genetic linkage analyses pinpoint the plausible cause to a recessive mutation in the 3’UTR of *Enox1* (Landouré et al. 2012). This mutation reduces *Enox1* transcript level to 20% of the normal level in B lymphoblastoid cell lines derived from homozygous patients, and 50% in those derived from healthy heterozygotes. Therefore, a hypomorphic *Enox1* mutation could cause MG, and transcriptome analyses in patient derived cells would determine whether they suffer splicing abnormalities. However, we noticed that some of the Italian patients lack autoantibodies against the neuromuscular junction component of acetylcholine receptor (Bergoffen et al. 1994). It is therefore possible that the underlying cause of MG in these patients is splicing defects in genes encoding important components of the neuromuscular junction, but not in autoimmunity exclusively as suspected. *Drosophila* lacks an adaptive immune function yet *Drosophila anw* mutants suffer muscle dysfunction, which might be similar in origin to ones in the Italian patients. Whether the development of mammalian neuromuscular junction is most sensitive to the loss of Anw/Enox1 require the establishment of mammalian animal models.

## Materials and Methods

### Drosophila crosses

#### General stocks

Fly stocks and crosses were cultured at 25°C on standard cornmeal food. Unless otherwise specified, *w^1118^* served as the wildtype stock. The *Df(3L)ED4486* stock (BL#8072), which has a deficiency of the *anw* region at 69E, was obtained from the Bloomington Drosophila Stock Center. A *mhc*-Gal4 stock was a kind gift from Dr. Zhuohua Zhang at Central South University of China.

#### CRISPR/Cas-9 mediated mutagenesis of anw

The sgRNAs used have the following sequences with the PAM sequence in bold: sgRNA#1, 5’-CCGGAAAATACAGATATCC**AGG**; sgRNA#2, 5’-GCCGCTAGTCACAAGAAGGTT**TGG**; sgRNA#3, 5’-GAGCGGGACTACGATAGACG **CGG**. All mutant alleles were verified by PCR followed by sequencing of homozygous adults.

#### Transgenic constructs and transgenic lines

All *anw* transgenic constructs were based on a 7.6kb genomic fragment of *anw* with the following genomic coordinates, 3L:12,843,772 to 12,851,417 (from release dm6, flybase.net), first subcloned into the vector pUASTattB. This fragment contains the *anw* coding region and approximately 500bp up- and 1300bp down-stream of the coding region. The *anw^GFP^* construct was made by inserting the coding region of GFP just upstream of the STOP codon of *anw*, using the method of bacterial recombineering (Zhang et al. 2014). Based on this GFP-containing construct, various *anw*-mutant constructs were made by PCR amplification of overlapping fragments followed by homology-directed assembly *in vitro*. These mutations are as follows with the genomic coordinates of the deleted sequence in parentheses: ΔRRM (3L: 12845392 to 12845574); Δα1 (3L: 12845629 to 12845706); Δα2/3 (3L: 12845743 to 12845844); Δα4 (3L: 12845863 to 12845955); Δα6 (3L: 12848730 to 12848966); Δc1 (3L: 12849462 to 12849674). The *gTc* mutation was constructed by site-directed mutagenesis changing the sequence of GGG to CCC (3L:12847263-5) and GG to CC (3L: 12847305-6). All constructs were verified by sequencing. All constructs were inserted at position 86F on chromosome 3R via phiC31-mediated site-specific integration, and they were combined with the endogenous *anw* mutant alleles via meiotic recombination. Primer sequences are provided in Table S2.

### Antibodies

Polyclonal antibodies against Anw were raised in mice against an N terminal (residues 1 to 437) and a C terminal (residues 439 to 804) antigen purified as inclusion bodies from *E. coli*. Sera were used in Western blots at 1:1000 dilution, and in immunostaining of muscle at 1:250. Commercial antibodies against human Enox1 (PA5-58355) and Enox2 (10423-1-AP) were purchased from ThermoFisher, and used at 1:5000 for Western blots. Anti-FLAG antibody (HT-201) was purchased from TransGen Biotech (China), and used at a dilution of 1:10000 on Western blots.

### Microscopic analyses of adult muscle structures and of protein localization

Muscle from adult thoraxes were dissected in cold PBS, fixed for 30 minutes in a fixative of 3.7% formaldehyde in PBS, washed three times for 15 minutes each with PBST (PBS plus 0.1% Triton T-100), stained with Phalloidin-TRITC (1:10000) from Sigma-Aldrich, and observed with a Zeiss LSM880 confocal microscope.

Various tissues were dissected in cold PBS, fixed in 4% formaldehyde in PBS for 20 minutes, washed in PBST three times. The fixed tissues were either stained with DAPI and observed directly for GFP fluorescence, or subjected to anti-Anw antibody staining followed by DAPI staining. Slides were observed on a Zeiss LSM880.

### Muscle protein enrichment

The back muscle of *Drosophila* adults was subject to three rounds of protein extraction, with the first two rounds aimed at removing muscle cell membrane proteins, while the final round yielding largely myofibrillar proteins. Thoraxes from 30 adults of mixed sexes were dissected in cold PBS, and homogenized with 200μl of extraction buffer 1 (150 mM NaCl, 0.5% NP-40) with a cryogenic grinder. After a 30 minutes incubation on ice, the sample was centrifuged at 13,000G for 10 minutes at 4°C, and the supernatant was collected as “extract #1”. Another round of extraction was performed identically as the first round except with a new extraction buffer (250 mM NaCl, 1% NP-40). The supernatant from this extraction does not contain much protein. A 200μl of an extraction buffer 2 (1 mM EDTA, 1% SDS) was used for the final round of extraction yielding “extract #2”.

### The minigene assay

#### Construction of expression plasmids

The target genomic fragments of the genes *bt* (4: 735910 to 737764 in release dm6) and *clc-a* (3R: 11798000 to 11799097) were amplified by PCR and subcloned into the expression vector pUASTattB. Deletion constructs of both *bt* and *clc-a* minigenes were constructed by homology-directed assembly *in vitro* and introduced into pUASTattB. The Gal4 coding region was amplified from genomic DNA from Gal4 expressing flies and cloned into the vector pAC5.1/V5-his (ThermoFisher) for Gal4 expression under the *actin5C* promoter. The *anw* cDNA isoform C and isoform B (flybase.net) were amplified from cDNA synthesized from wildtype total RNA, and subcloned into pUASTattB for UAS-controlled expression of Anw. Isoform B potentially encodes a 50% shortened and non-functional Anw protein, and was used as a negative control in the minigene assay.

Full length cDNA clones of human *Enox1* and *Enox2* were recovered by RT-PCR amplification from total RNA of human 293T cells, and subcloned into pUASTattB for expression in *Drosophila* cells. cDNAs of Aplysia *anw* and zebra fish *Enox1* and *Enox2* were synthesized by Sangon Biotech company of China, and subcloned into pUASTattB. They were also tagged at the C-terminus with 3XFLAG to allow protein detection on Western blots.

Expression plasmids for the series of *anw* deletion mutations were constructed by PCR amplification of overlapping fragments followed by *in vitro* homology-directed assembly. All constructs were verified by sequencing. Primer sequences are provided in Table S2.

#### S2 Cell culturing and transfection

S2 cells were maintained at 25°C with InsectPro® Sf9 medium supplemented with 10% heat-inactivated FBS and antibiotics. Cells were seeded into a 6-well plate at a density of 5×10^5^ cells/mL about 6 hours before transfection, and the culture normally reached about 70% confluency at the time of transfection. Transfection was performed using FuGENE® HD Transfection Reagent, according to the manufacturer’s instructions. For a 6-well plate, a ratio of 1μg of total plasmid DNA and 3μl of transfection reagent in InsectPro® Sf9 medium was followed for each transfection. Transfected cells were harvested and analyzed 48 hours later.

### RNA purification, RT-PCR, qPCR and Immunoprecipitation

Total RNA was extracted from adults, dissected tissues, or S2 cells using TRIzol reagent (Ambion, 15596018) according to the manufacturer’s instructions. For RT-PCR and qPCR, cDNA was synthesized using the PrimeScript RT reagent Kit (TaKaRa, RR047A).

For RNA IP, we followed the protocol by Gagliardi and Matarazzo (2016). Briefly, cells were suspended in lysis buffer (100 mM KCl, 5 mM MgCl_2_, 10 mM HEPES-NaOH pH 7, 0.5 % Nonidet P-40, 1 mM DTT, 200 units/ml of RNase OUT, and EDTA-free Protease Inhibitor Cocktail), followed by a freeze-thaw treatment for three times, with each time lasting 10 minutes. The supernatant from a 15 min centrifugation at 16000G was incubated for 3 hours at 4°C with primary antibodies followed by precipitation with protein A/G agarose beads. RNA purified from the beads was used in RT-PCR, qPCR or next-generation sequencing. Quantitative RT-PCR reactions for Figure S5 were carried out on a Real-time PCR machine (QuantStudio 5 by Applied Biosystems), according to a protocol from the KAPA SYBR FAST qPCR Kit (KK4601). Three biological replicates were tested for each sample. The percent over input was calculated according to the protocol above and presented as the bar graph in Figure S5. Primer sequences are provided in Table S2.

### RNA sequence and RNA IP sequence analysis

We performed whole transcriptome sequencing (RNA-seq) of RNA samples recovered from wildtype adults of mixed sexes and dissected thorax samples from adults of mixed sexes. The genotypes were *w^1118^* as wildtype and *anw^1/3^* as mutants.

The Novogene company (China) performed transcriptome sequencing. Briefly, mRNA was purified from total RNA using poly-T oligo-attached magnetic beads. The constructed library was purified (AMPure XP system), and library quality was assessed on the Agilent Bioanalyzer 2100 system. The library was sequenced on an Illumina Novaseq platform, generating 150 bp paired end reads.

The raw data undergoes processing with Trim Galore (0.6.10) to remove adapters and low-quality reads with the following parameters: --paired –quality 20 –length 30. For genome alignment, STAR (2.7.1a) generates an index with these parameters: --runThreadN 8, --runMode genomeGenerate, and --sjdbOverhang 149. Then, the sequences are aligned to the Drosophila genome. Subsequently, splicing analysis is conducted using rMATS (4.2.0) on the cleaned reads with the following parameters: -t paired, --readLength 150, and --nthread 4. Both rMATS and STAR utilize Ensembl release 109 as the reference genome. The tables of significant alternative splicing events are filtered by FDR < 0.01 and Difference in inclusion levels > 5%.

### Homology searches

To search for Anw homologous proteins, a BLAST search was first performed with either the *Drosophila* or the human proteins excluding the RRM domain. Once putative homologs were identified, the presence of an annotated RRM domain at the N-terminus was used to definitively identify Anw homologs. To search for the conserved small exon in *anw* homologous genes, the amino acid residues encoded by the exon and its flanking ones were used to identify genomic sequences of the region, and exon/intron structures were then identified manually.

## Acknowledgments

We thank Mr. Zheng Ou and Dr. Yang Shen at the University of South China for technical assistance, Drs. Zhuohua Zhang and Zhonghua Hu from Central South University for sharing cell lines and reagents, Dr. Weichao Zhao from USC for sharing zebra fish adults, and Drs. Yujie Fan and Yongzheng Xu at the Wuhan University, and Dr. Natalia Misunou at EMBL for critical comments on the manuscript. for critical comments on the manuscript. We thank Ms. Liqian Chen at Sun Yat-sen University for her assistance in the initial stage of the project.

## Author Contributions

Conceptualization: TZ, KL, YX, YSR Methodology: TZ, KL, YSR Investigation: TZ, YX, XHT, XYT, YSR Visualization: TZ, KL, YX, MC, YSR Supervision: YSR Writing—original draft: TZ, KL, MC, YSR Writing—review & editing: TZ, KL, YX, XYT, MC, YSR

## Data and Materials Availability

All data needed to evaluate the conclusions in the paper are present in the paper and/or the Supplementary Materials. *Drosophila* stocks, plasmids and antibodies are freely available from YSR.

## Supplemental Materials

Supplemental Result 1. Additional splicing events affected in *anw* mutants shown in Figure S3

### *zasp67*, exons 6-7-8-9-10

In the wildtype sample, the joining of exons 6, 7 and 10 predominates (“b” in Figure S3). In the absence of Anw, AS events utilizing exons 8 and 9 increases even ones without the previously dominant exon 7. Therefore, Anw appears to promote the exclusion of specific exon pairs for *zasp67* splicing.

### *CG5707,* exons 1-2-3

There appears to be an increase of the inclusion of exon 2 in *anw*-mutants. This was verified by sequencing of the products indicated as “inclusion” in Figure S3.

### *teq,* exons 1-2

There appears to be a decrease of the spliced product leading to a lengthening of the 5’UTR in the mutants.

### *nrm,* exons 20-21-22-23-24

There appears to be a decrease in the “spliced” product (“a” in Figure S3) and concomitant increase in other larger spliced products such as “b” in Figure S3.

### atypical protein kinase C (aPKC), exons 11-12-13-14

Exons 12 and 13 are mutually exclusive. In *anw* mutants, the frequency of exon 12 selection was reduced accompanied by intron retention between exons 11 and 14.

### *csp,* exons 4-5-6

The “skipped” product of exon 5 is decreased in the mutants.

### *norpA,* exons 3-4-5-6

In the mutant, the inclusion of exon 4 is markedly increased.

### *CG43897,* exons 1-2-3-4-5

The inclusion of exons 3, 4, and 5 is through selection of the different 3’ splice sites. The *anw* mutations affected this selection in a complex pattern. Nevertheless, the decrease in products “b” and “c” is evident in the mutants.

### CG43427 5’ UTR

The very 5’ of the *UTR* has a series of AS events utilizing different 5’ ss but sharing the same 3’ ss. In wildtype adults, a particular 5’ss is allowed to be utilized whereas its uses are inhibited in the mutants.

### Fer1HCH 5’ UTR

The selection of different 3’ splice sites determines the length of the 5’ *UTR* and the *anw* mutation affects this selection. Specifically, product “c” is markedly reduced in the mutants.

### 5-HT2A 5’UTR

The appearance of ectopic spliced products is apparent in the mutant samples (* in Figure S3). Sequencing of this product indicates that it is derived from this region but contains 21nt additional sequences which origin we could not currently explained.

## Supplemental Result 2. Anw functions by interacting with its RNA targets as shown in Figure S5

We modified the Anw-expressing vector to include a C-terminal 3xFLAG tag and confirmed the functionality of the Anw-3XFLAG protein to direct normal splicing of the minigenes (Figure S6). The transfected cells were split in half, one half was subjected to anti-FLAG IP to isolate Anw-RNPs. The other half was used as a control where unrelated IgG was used for the pull-down. This approach eliminated a potential complication of differences in transfection efficiency between cell populations. We nevertheless carried out a modified version of this approach, in which two cell populations were transfected with untagged or FLAG-tagged Anw expression plasmids followed by anti-FLAG pull-down, and we obtained very similar results (see the “third approach” below). We first used regular RT-PCR to investigate Anw-RNA interaction in a semi-quantitative way. As shown in Figure S5A, we designed primers to detect both spliced and unspliced products across multiple introns in the minigenes, and included a regular PCR using genomic DNA as a template to mark the running positions of unspliced products. As a control, we measured the endogenous *rp49* levels in both the input and IP samples. As an additional control, we performed the same series of RNA-IP experiments with a minigene from a region of the *agps* gene, which is not under the regulation of Anw in animals. It is clear from Figure S5A that our Anw-FLAG protein was able to specifically enrich pre-mRNA fragments from both the *bt* and *clc-a* minigenes, supporting our model in which Anw interacts with its target as an RNA binding protein. We used RT-qPCR and provided a more quantitative confirmation of the interaction between Anw and its RNA targets (bar graph in Figure S5A). In a third approach, we FLAG-IPed two populations of cells, one expressing Anw and one Anw-FLAG, and both expressing the *clc-a* minigene. After re-confirming by RT-PCR that Anw-FLAG could specifically pull down *clc-a* pre-mRNA (Figure S5B), we sent an RNA-IP sample from this experiment for sequencing. As shown in the genome browser snapshots in Figure S5B, the entire region of the expressed fragment is enriched with RNA-seq reads as compared with the reads from control IP. This again confirms the interaction between Anw and the pre-mRNA of *clc-a*. We were hoping that RNA-seq could reveal distinct peaks of enrichment along the pre-mRNA fragment hence identifying high affinity sites of Anw binding. However, the RNA-seq results suggest that Anw binds the entire pre-mRNA target, consistent with earlier RT-PCR and mini-gene deletion results (Figure 4). We wish to point out that this series of RNA-IP experiments differ from the state-of-the-art CLIP method in that UV crosslinking or target trimming by RNase treatment was not implemented. These critical differences might have reduced the resolution below what is needed for mapping Anw binding sites. Nevertheless, multiple approaches with the minigene system support a direct interaction of Anw with its RNA targets, even though we have not succeeded in identifying any candidate motifs that Anw binds.

## Supplemental Result 3. The effects of Anw mutations on minigene splicing in S2 cells as shown in Figure S6

To take advantage of the minigene assay established above, we constructed six expression vectors each containing an in-frame deletion disrupting a potentially structured domain, and two vectors deleting the disordered C-terminal regions (Figure S6A). In the last and ninth vector, we constructed a point mutation within the short helix 5 (Figure S6B) and named it *anw^gTc^*, which we will discuss in the main text. These nine vectors were transfected into S2 cells individually, together with either the *bt* or the *clc-a* minigene. RT-PCR was used to assay the impact of Anw truncations on splicing. As shown in Figure S6C, internal deletions of the structured elements of Anw all negatively affect splicing. Remarkably, deletions of the C terminus, even the C2 deletion of 313 amino acids (over a third of the protein) did not significantly affect minigene splicing. We refrained from drawing conclusions comparing the effect of different mutants considering the potential issue of expression level differences as a result of experimental variations and/or stability of the target proteins. Indeed, Western blotting detected a large variability in Anw levels (Figure S6C). Nevertheless, we have identified potentially important domains for Anw function using the minigene assay.

---

**Figure S1.**
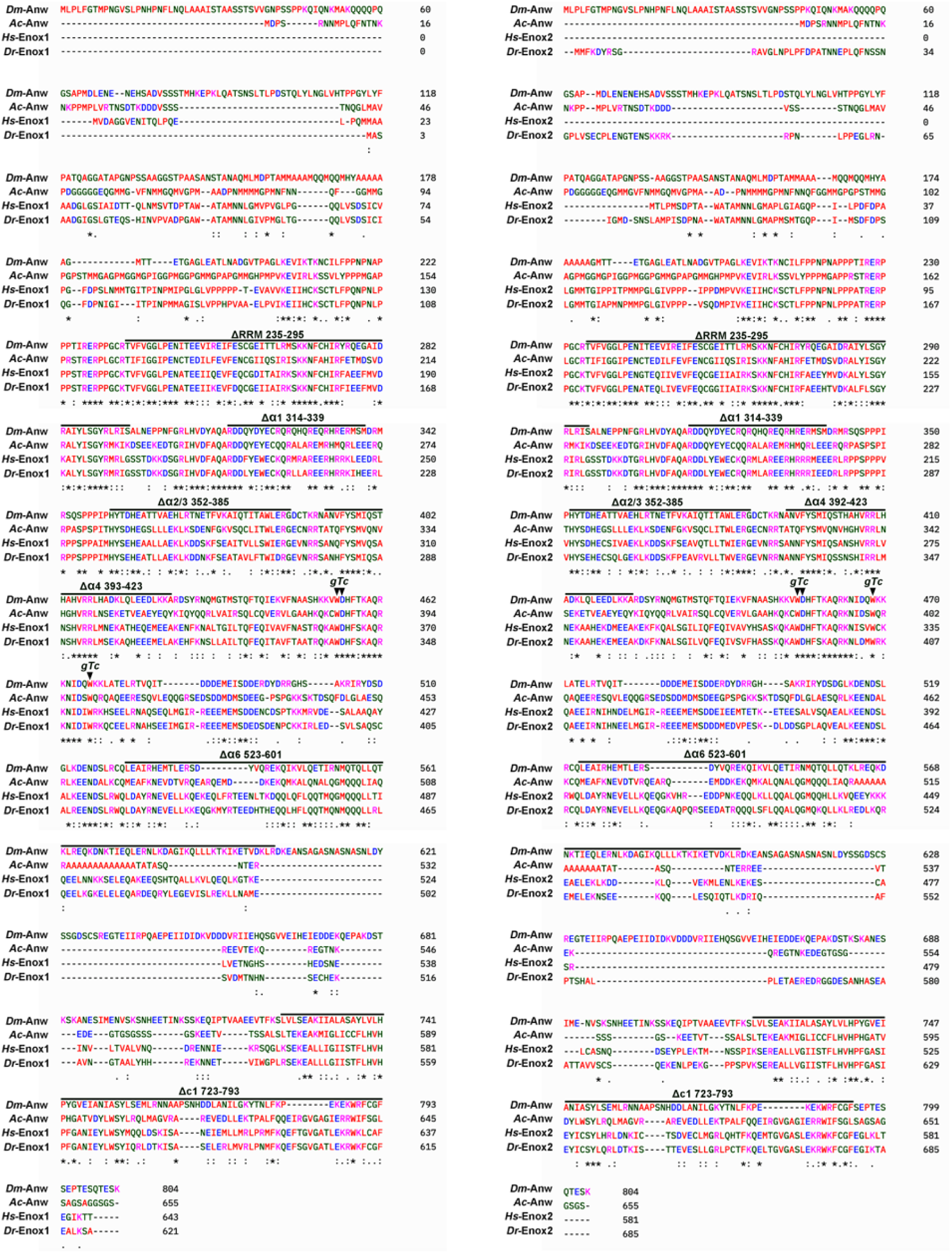
Anw homologous proteins. A clustal alignment of Anw homologous proteins from *Drosophila melanogaster* (Dm), *Aplysia californica* (Ac), *Homo sapiens* (Hs) and *Danio rerio* (Dr) is given separately for Anw *versus* Enox1, and Anw *versus* Enox2. A series of transgenes with internal deletions were constructed to identify functionally important domains of Anw (see main text). The deleted regions are marked with horizontal lines with the names and the deleted residues given on top. The three residues changed in the *anw^gTc^* mutation are marked with arrowheads.

**Figure S2.**
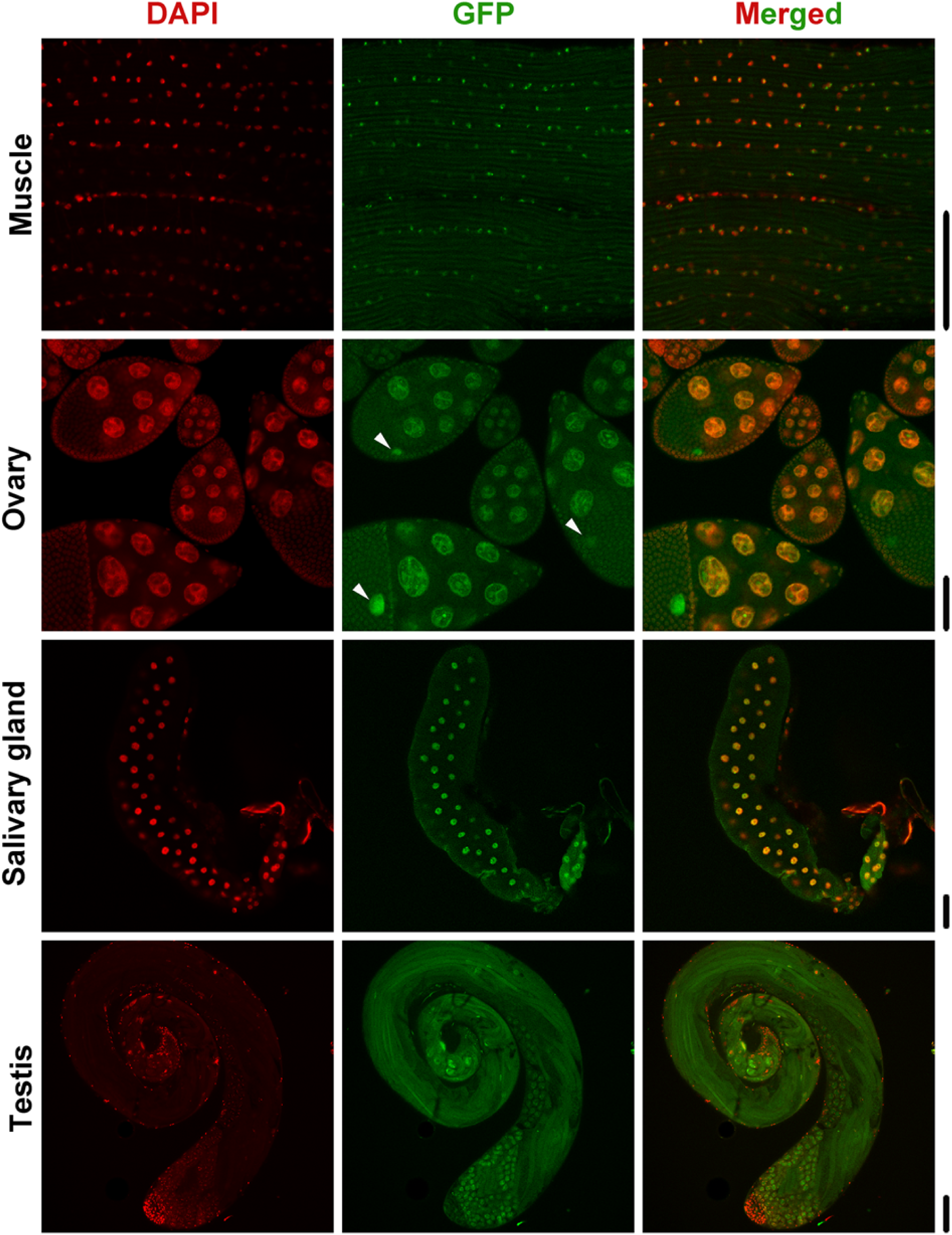
Anw localization in different tissues. A supplemental to Figure 2 showing lower magnification images of Anw^GFP^ signals in different tissues. The genotype is *anw^1^, anw^GFP^*. Arrowheads indicate the position of the nucleus of the oocyte. Scale bars are 80μm.

**Figure S3.**
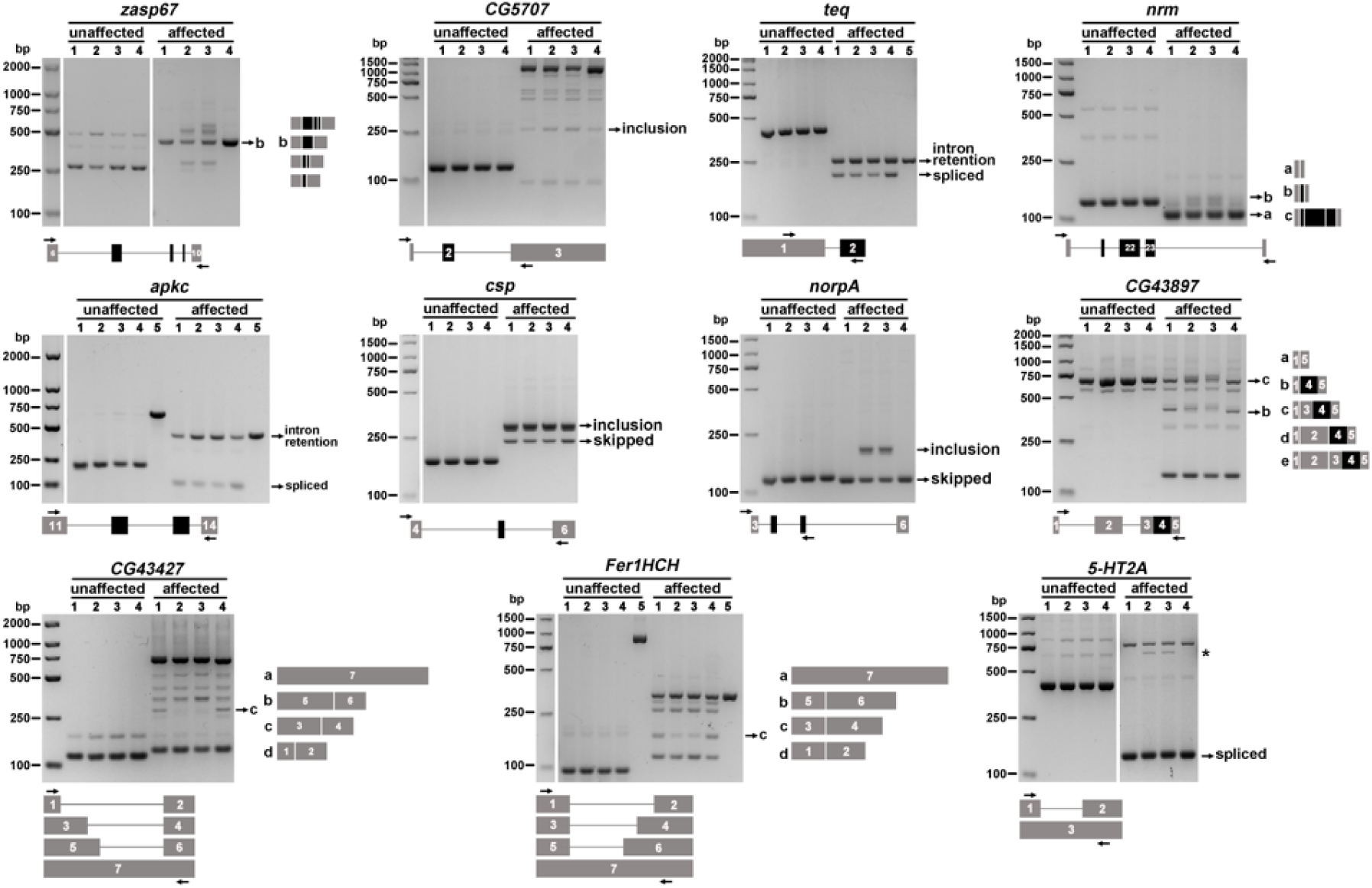
Additional splicing defective events in *anw* mutants. This is supplemental to Figure 3. For each gene a gel picture showing results from two RT-PCR reactions using total RNA from adults, with one primer pair spanning a region not controlled by Anw (“unaffected”). For their approximate positions see Figure S4. Another pair of primers spans the “affected” region, which are indicated as arrows in the diagram of the exon/intron structure shown under the gel picture with exon labelling according to that in Figure S4. The products on the gel are either labelled with a description: “spliced”, “skipped”, “intron retention”, “inclusion”, or a diagram illustrating their structure labelled with a, b, c *etc*. The genotypes for the lanes are as followed and indicated in parentheses: 1 (*wt*); 2 (*anw^1/1^*); 3 (*anw^3/df^*); 4 (*anw^1^, anw^GFP^*). For lane 5, genomic DNA from *wt* adults was used as the PCR template to indicate the running position of the un-spliced products.

**Figure S4.**
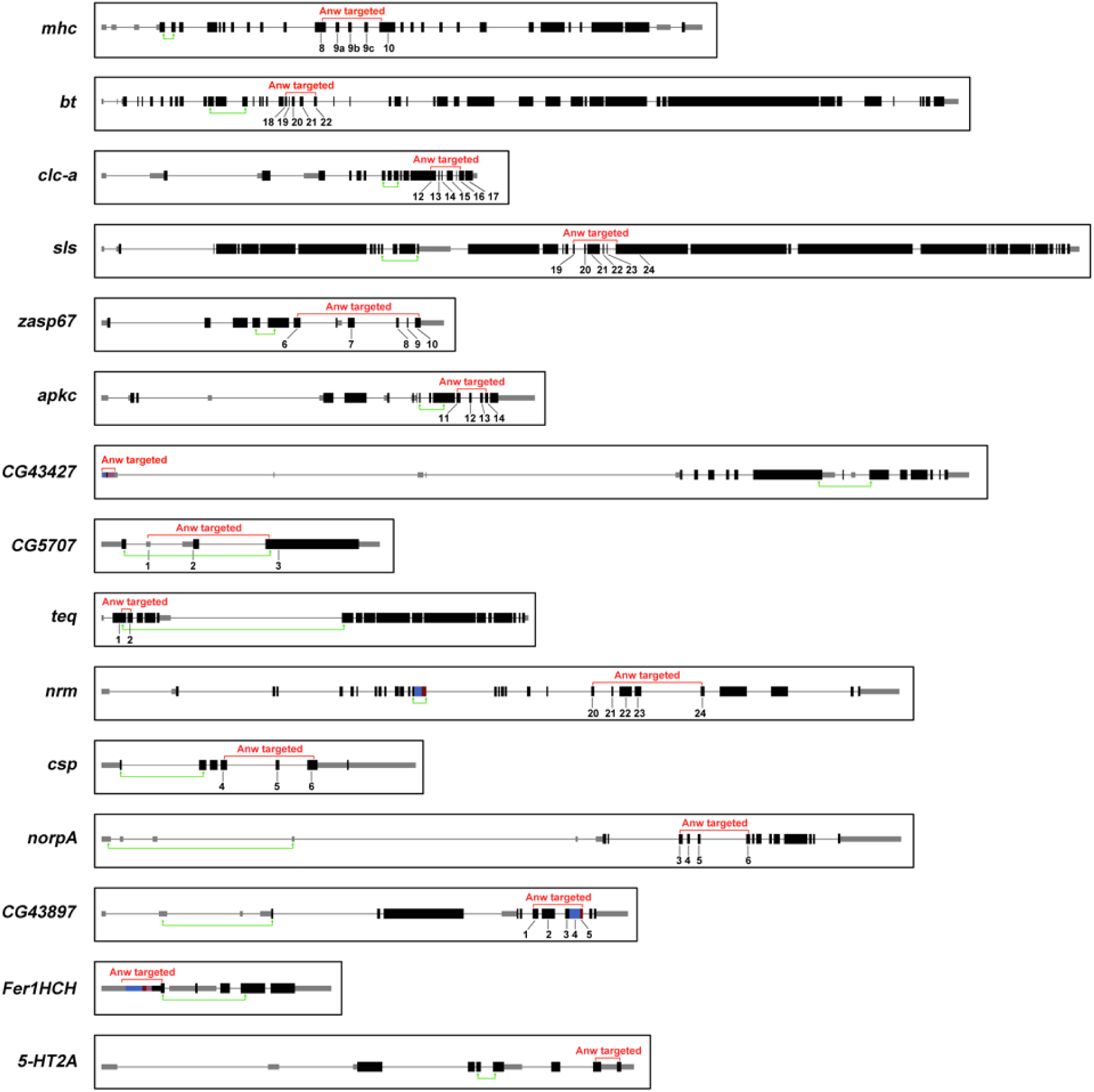
Gene structure and splicing programs of *anw*-mutant affected loci. The exon/intron structures of 15 *anw*-controlled loci are shown with boxes representing exons and lines representing introns. Black labelling indicates coding exons and grey one UTRs. Two regions of each gene were chosen for RT-PCR analyses. The “Anw targeted” region (in red) is marked on top of the diagram with the affected exons numbered underneath. The “unaffected” region (in green) is marked under the diagram with two vertical arrows.

**Figure S5.**
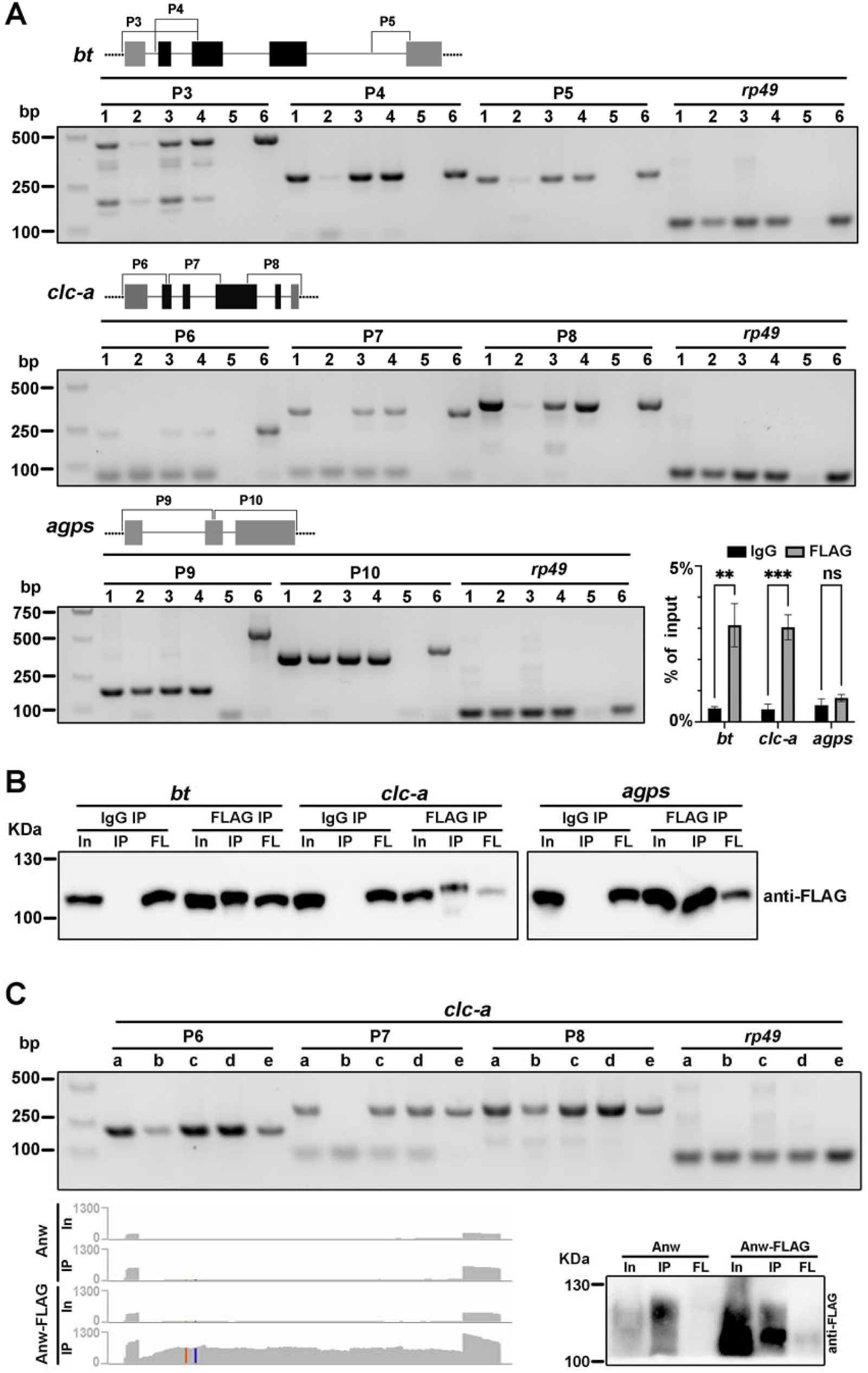
Anw interacts with its RNA targets A. Investigating Anw-RNA interaction by RNA-IP. For each minigene (three in total), a gel picture of a semi-quantitative RT-PCR reaction is shown under the diagram of the exon/intron structure of the minigene along with designations of the different PCR products (P3-P10). Dotted lines represent regions of the vector flanking the minigene. Cells transfected with the minigene reporters along with a plasmid expressing Anw-FLAG protein was used in IP. Lane 1: input from IgG IP; lane 2: IP from IgG IP; lane 3: input from FLAG IP; lane 4: IP from FLAG IP; lane 5: IP from FLAG IP with reverse transcriptase omitted in the RT reaction; lane 6: regular PCR templated off of genomic DNA for identification of unspliced products. A primer pair for *rp49* was included as a control in all RT-PCR reactions. For *clc-a*, smaller (<100bp) and nonspecific products are evident. An RT-qPCR based quantification is given in the bar chart. ns: not significant; **: p<0.01; ***: p<0.001. **B. Western blotting versification of FLAG-IP effectiveness**. For each pair of RNA-IP experiments (IgG and FLAG), extracts were run and blotted with anti-FLAG. **C. Further investigation of Anw-RNA interaction by RNA-IP**. A gel picture of a semi-quantitative RT-PCR reaction is shown. Cells transfected with the minigene along with a plasmid expressing the full length Anw protein (lanes a and b) or the Anw-FLAG protein (lanes c and d) were used in RNA-IP. Lanes a and c are PCR with input RNA, and lanes b and d are PCR with IP RNA. Lane e is a regular PCR templated off of genomic DNA for the identification of unspliced products. *rp49* was used as a control. A diagram of the *clc-a* minigene is shown to the right of the gel picture with the positions of the primer pairs for detecting IP products shown. Underneath this diagram is an IGV screen shot of the minigene region where RNA seq reads from input and IP were loaded. Next to the screen shot shows an anti-FLAG Western blot confirming the effectiveness of the FLAG-IP. In: input extracts; IP: precipitated proteins; FL: flowthrough extracts after IP.

**Figure S6.**
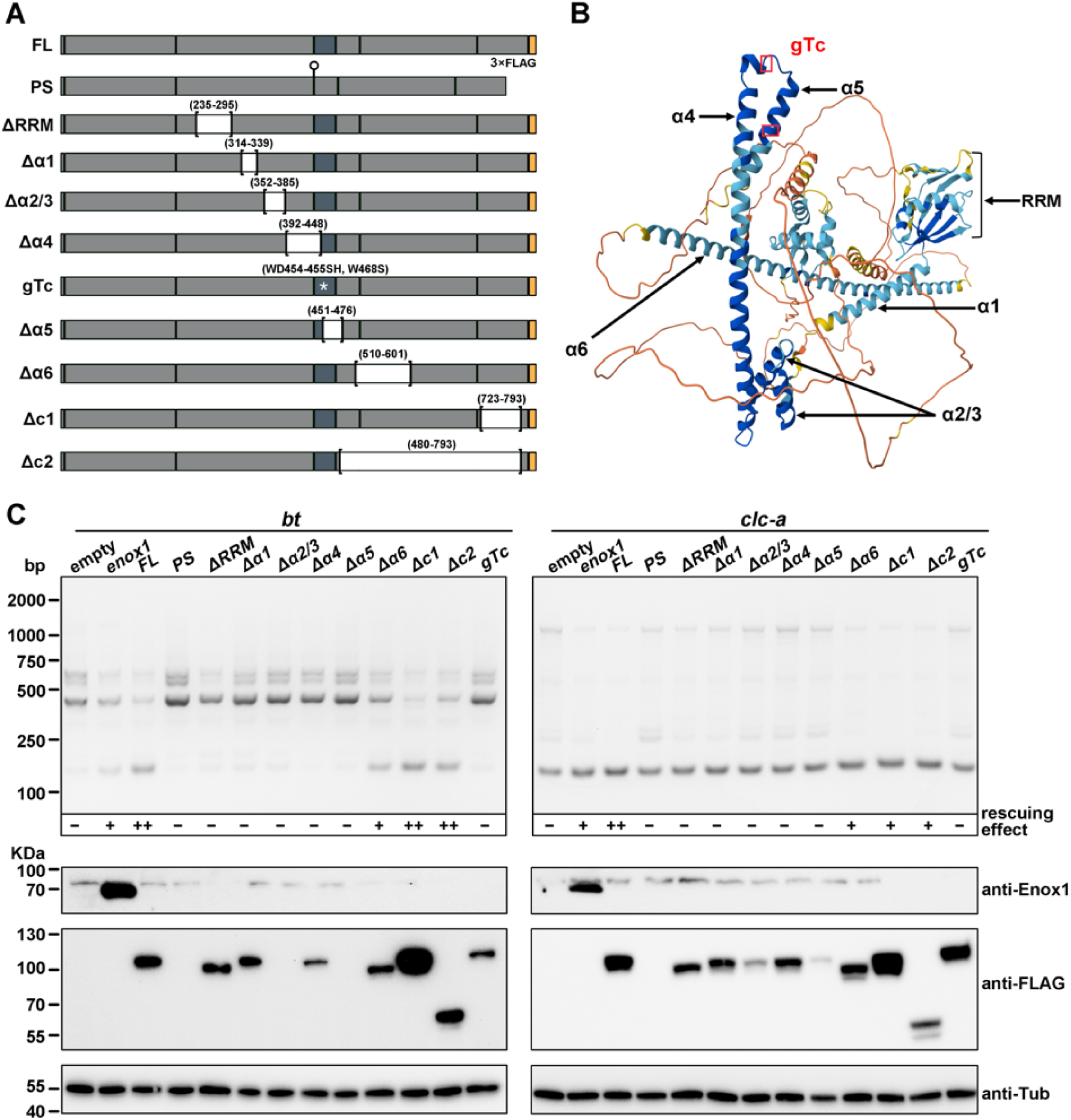
Structural domains of Anw and the effects on mini-gene splicing upon their disruption. **A. Domain deletion mutations of *anw***. A schematic of Anw and its derivative proteins are shown with boxes representing coding regions in which the exon/intron junctions are marked with vertical lines. “FL” represents full-length Anw protein. In the internally deleted Anw proteins, the deleted regions are demarcated with brackets and labelled white. The deleted residue numbers are listed on top in parentheses. All Anw proteins are tagged at the C-terminus with 3XFLAG (orange box) except the one with a premature STOP codon (PS, marked with a lollipop) due to the exclusion of a small exon (labelled dark grey). The position of the *anw^gTc^* mutation is marked with a white star. **B. Anw structure by Alpha-fold**. Each structured element is given a designation and marked with arrows. The position of the two changes in the *anw^gTc^* mutation is marked with red boxes. **C. The effects of Anw mutations on minigene splicing**. The top panels are gel pictures showing RT-PCR detection of splicing products from the *bt* (left) and *clc-a* (right) minigenes. A score for “rescuing effect” is given for each mutation. The Anw proteins expressed along with the minigenes are listed on top of the gel pictures. “empty” indicates that the empty expression vector was transfected. “enox1” indicates that the human Enox1 expressing vector was transfected. The other designations are the same as shown in **A**. Under the gel pictures are Western blot results accessing the expression of the Anw proteins. Except for human Enox1 expression, all the other Anw proteins are detected with anti-FLAG.

**Figure S7.**
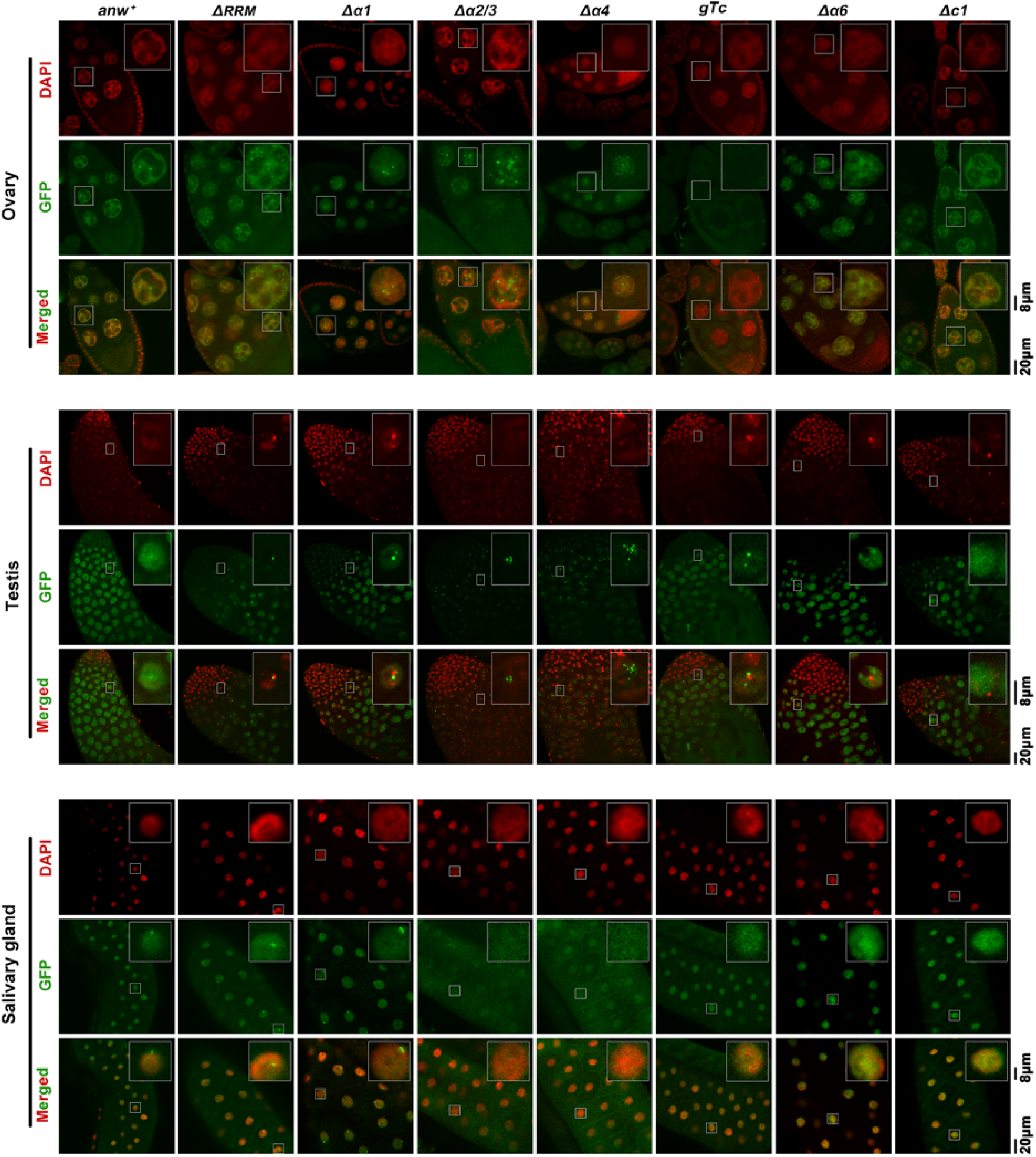
Domains of Anw important for nuclear body formation in different tissues. This is supplemental to Figure 6. GFP florescence is used to assess the effect of Anw mutations on its localization in adult ovaries, testes and larval salivary glands. In each panel, DAPI signals are in red, GFP in green. A single nucleus is chosen for showing a magnified image.

**Video S1. *anw^1^* flies are flightless**

Video shows wildtype flies (in the left vial) fly or jump after being disturbed while *anw^1^*flies (in the right vial) lose those abilities.

**File S1. AS events identified in thorax RNA-seq**

Bioinformatic analyses of AS changes from RNA-seq of thorax samples. Six tables are included showing individual analysis for A3SS, A5SS, MXE, RI, SE and their combined data.

**File S2. AS events identified in whole-fly RNA-seq**

Bioinformatic analyses of AS changes from RNA-seq of whole-fly samples. Six tables are included showing individual analysis for A3SS, A5SS, MXE, RI, SE and their combined data.

